# Natural variation in yeast reveals multiple paths for acquiring higher stress resistance

**DOI:** 10.1101/2023.11.20.567940

**Authors:** Amanda N. Scholes, Tara N. Stuecker, Stephanie E. Hood, Cader J. Locke, Carson L. Stacy, Qingyang Zhang, Jeffrey A. Lewis

## Abstract

**Background:** Organisms frequently experience environmental stresses that occur in predictable patterns and combinations. For wild *Saccharomyces cerevisiae* yeast growing in natural environments, cells may experience high osmotic stress when they first enter broken fruit, followed by high ethanol levels during fermentation, and then finally high levels of oxidative stress resulting from respiration of ethanol. Yeast have adapted to these patterns by evolving sophisticated “cross protection” mechanisms, where mild ‘primary’ doses of one stress can enhance tolerance to severe doses of a different ‘secondary’ stress. For example, in many yeast strains, mild osmotic or mild ethanol stresses cross protect against severe oxidative stress, which likely reflects an anticipatory response important for high fitness in nature.

**Results:** During the course of genetic mapping studies aimed at understanding the mechanisms underlying natural variation in ethanol-induced cross protection against H2O2, we found that a key H2O2 scavenging enzyme, cytosolic catalase T (Ctt1p), was absolutely essential for cross protection in a wild oak strain. This suggested the absence of other compensatory mechanisms for acquiring H2O2 resistance in that strain background under those conditions. In this study, we found surprising heterogeneity across diverse yeast strains in whether *CTT1* function was fully necessary for acquired H2O2 resistance. Some strains exhibited partial dispensability of *CTT1* when ethanol and/or salt were used as mild stressors, suggesting that compensatory peroxidases may play a role in acquired stress resistance in certain genetic backgrounds. We leveraged global transcriptional responses to ethanol and salt stresses in strains with different levels of *CTT1* dispensability, allowing us to identify possible regulators of these alternative peroxidases and acquired stress resistance in general.

**Conclusions:** Ultimately, this study highlights how superficially similar traits can have different underlying molecular foundations and provides a framework for understanding the diversity and regulation of stress defense mechanisms.

**Author Summary:** Organisms in nature frequently experience environmental stress in predictable patterns. For example, during the summer months in temperate climates, the warmth of the morning sun gradually gives way to high afternoon temperatures. Organisms that can anticipate these predictable patterns to mobilize stress defenses would have an advantage in nature. One way organisms anticipate future stress is through ‘cross protection,’ where cells exposed to a mild dose of one stress gain the ability to survive an otherwise lethal dose of different stress. To better understand the molecular mechanisms that are responsible cross protection, we have been taking advantage of wild yeast strains that are either more resilient or more sensitive to stresses. During the course of this study, we found that strains with superficially similar levels of cross protection differ in the precise molecular mechanisms that underlie the trait. Our study suggests that different molecular strategies may be important for yielding similar stress resistances under different environmental constraints, and highlights the power of harnessing natural genetic diversity to understand the molecular mechanisms underlying differences in environmental responses.

## Introduction

In nature, environmental stress often occurs in predictable patterns. Extremes of temperature and precipitation follow seasonal patterns, and stress gradients can also predictably occur over even shorter timescales. During the daytime, temperature predictably increases, and while in estuary environments, salinity predictably increases closer to seawater sources [1]. In addition to single-stress gradients, some stresses tend to co-occur or happen in predictable succession. For example, high temperatures frequently coincide with low precipitation [2], so organisms often experience both heat and drought stresses simultaneously [3].

Because stresses can occur in predictable patterns, organisms that can anticipate these patterns would likely have a fitness advantage in nature. One such anticipatory strategy is a phenomenon called acquired stress resistance, where exposure to a mild dose of stress can enable organisms to survive an otherwise lethal dose of severe stress. Acquired stress resistance can occur when the mild and severe stresses are the same (same-stress protection) or are different (cross protection). Cross protection has been observed in diverse organisms ranging from prokaryotes to humans under a wide variety of circumstances. For example, mild drought pretreatment protects several crop plants against severe heat stress [4–6], which has major implications for mitigating the effects of climate change. In terms of human health, fever temperatures protect against oxidative stress, which may be important for immune cells experiencing increased generation of reactive oxygen species during infection [7], and transient whole-body hyperthermia (i.e., a mild heat shock) cross protects against oxidative damage caused by reperfusion of oxygenated blood following surgery [8, 9]. Therefore, understanding the molecular mechanisms underlying acquired stress resistance and cross protection has wide-ranging applications.

While a large number of genetic screens in the budding yeast *Saccharomyces cerevisiae* have been performed to understand the genetic basis of the intrinsic resistance of unstressed cells [10–38], only a limited number of studies have investigated acquired stress resistance [39–43]. An emerging theme though in studies of acquired stress resistance is regulatory complexity and partial redundancy. When comparing genetic screens for acquired H2O2 resistance following different mild pretreatments (H2O2, salt, heat, and dithiothreitol) [39–41], the lack of overlap in identified regulatory mutants strongly suggests that each mild- pretreatment triggers specific regulators that induce high H2O2 resistance. This is evocative of “many-to-one mapping” seen in morphological evolution [44], where multiple combinations of different structural parts can lead to the same functional outcome. In this case, the “multiple means to the same end” proposed by Berry and colleagues [41] can include multiple signaling pathways converging on the same molecular solution for acquiring higher stress resistance. It is also possible that activation of many different antioxidant defenses can lead to similar H2O2 resistance outcomes. However, identification of the direct antioxidant defense genes necessary for acquired H2O2 resistance has had limited success [39–41], suggesting at least partial redundancy in the antioxidant defenses responsible for acquired H2O2 resistance.

Additionally, while cross protection is widespread in yeast, it is not universal for all stress combinations. Berry and colleagues [45] found that mild heat stress protected cells against all severe stresses tested (NaCl, H2O2, ethanol, and heat), while mild NaCl only protected against severe NaCl and severe H2O2. The observed patterns of stress cross protection likely mirror the sequential order in which organisms typically encounter stresses in their natural environments [46]. However, conflicting reports about which stresses can or cannot confer cross protection within a given species are common (reviewed in [47]). At least for the case of the yeast ethanol response, we have shown that cross protection phenotypes can differ dramatically depending on the genetic background of the strain [48–50]. While perhaps not surprising that the genetic context affects a complex trait such as cross protection, it does suggest that different genetically distinct populations of yeast may have different anticipatory responses that reflect specific selective pressures unique to their ecological niches.

Most of our limited knowledge about the mechanisms underlying acquired stress resistance in yeast is restricted to laboratory strains [39, 41, 42]. However, heterogeneity in acquired resistance phenotypes clearly occurs and can have broad impact. For example, cancer cells appear to have defective acquired stress resistance phenotypes, which can be targeted to enhance the effectiveness of chemotherapy while sparing healthy cells [51, 52]. We have been leveraging natural variation in acquired stress resistance phenotypes to better understand both the genetic architecture of the trait and especially the stress defense genes ultimately responsible for the phenotype.

We previously reported that our commonly-used laboratory yeast strain (S288c) fails to acquire further H2O2 resistance when pretreated with a mild dose of ethanol, while many wild yeast strains can [50]. To understand the genetic basis of this trait, we performed genetic mapping in a cross between the lab strain and a wild oak strain (YPS163). This analysis identified the causal polymorphism—a transposon insertion in the lab strain that partially inactivates the heme-activated transcription factor Hap1p, leading to decreased ethanol-responsive expression of a key H2O2 scavenging enzyme, cytosolic catalase T (Ctt1p) [50]. In that wild oak strain, a *ctt1Δ* mutation completely eliminated ethanol-induced cross protection against H2O2 [50], suggesting the absence of compensatory mechanisms for acquired H2O2 resistance under that condition.

In this study, we found surprising heterogeneity across diverse yeast strains in whether *CTT1* function was fully necessary for acquired H2O2 resistance. Some strains exhibited partial dispensability of *CTT1* when ethanol and/or salt were used as mild stressors, suggesting that compensatory peroxidases may play a role in acquired stress resistance in certain genetic backgrounds. We leveraged global transcriptional responses to ethanol and salt stresses in strains with different levels of *CTT1* dispensability, allowing us to identify possible regulators of these alternative peroxidases and acquired stress resistance in general. Ultimately, this study highlights how superficially similar traits can have different underlying molecular foundations and provides a framework for understanding diversity and regulation of stress defense mechanisms.

## Results

### Strain-specific catalase (Ctt1p) requirements for acquired oxidative stress resistance

While a commonly-used S288c-derived lab strain of yeast is unable to acquire further H2O2 resistance following mild ethanol pretreatment, most wild yeast strains can [50].

Experiments designed to identify the mechanistic basis of this difference led to the discovery that cytosolic catalase T (Ctt1p)—a major H2O2-scavenging enzyme—plays a key role.

Specifically, loss of *CTT1* function in a wild oak strain (YPS163) completely abolished ethanol-induced cross protection against H2O2 [50].

We initially suspected the involvement of *CTT1* in ethanol-induced cross protection because a previous study found that *CTT1* was necessary for salt-induced cross protection against H2O2 in the S288c lab strain [41]. Thus, when analyzing the phenotype of YPS163 *ctt1Δ* mutants for ethanol-induced cross protection, we included salt-induced cross protection as a “positive control.” Surprisingly, YPS163 *ctt1Δ* mutants retained substantial acquired H2O2 resistance following mild salt pretreatment (∼25% of wild-type resistance or an additional 1mM of resistance compared to the mock-treated control, Fig 1), a level of resistance expected to be physiologically relevant for fitness in nature [53, 54]. In contrast, and consistent with previous studies [41], no residual acquired resistance was detected in the S288c background following mild salt pretreatment. Consistent with our previous studies [50], we did observe that mild ethanol pretreatment renders S288c even more sensitive to H2O2. Surprisingly, deletion of *CTT1* in S288c eliminated the sensitizing effects of ethanol pretreatment on H2O2 resistance. Overall, the results from this experiment suggested that *CTT1* was conditionally necessary for acquired H2O2 resistance depending on the identity of the primary stress and the genetic background (i.e., a gene-environment (GxE) interaction), which we sought to explore further.

**Figure. 1.**
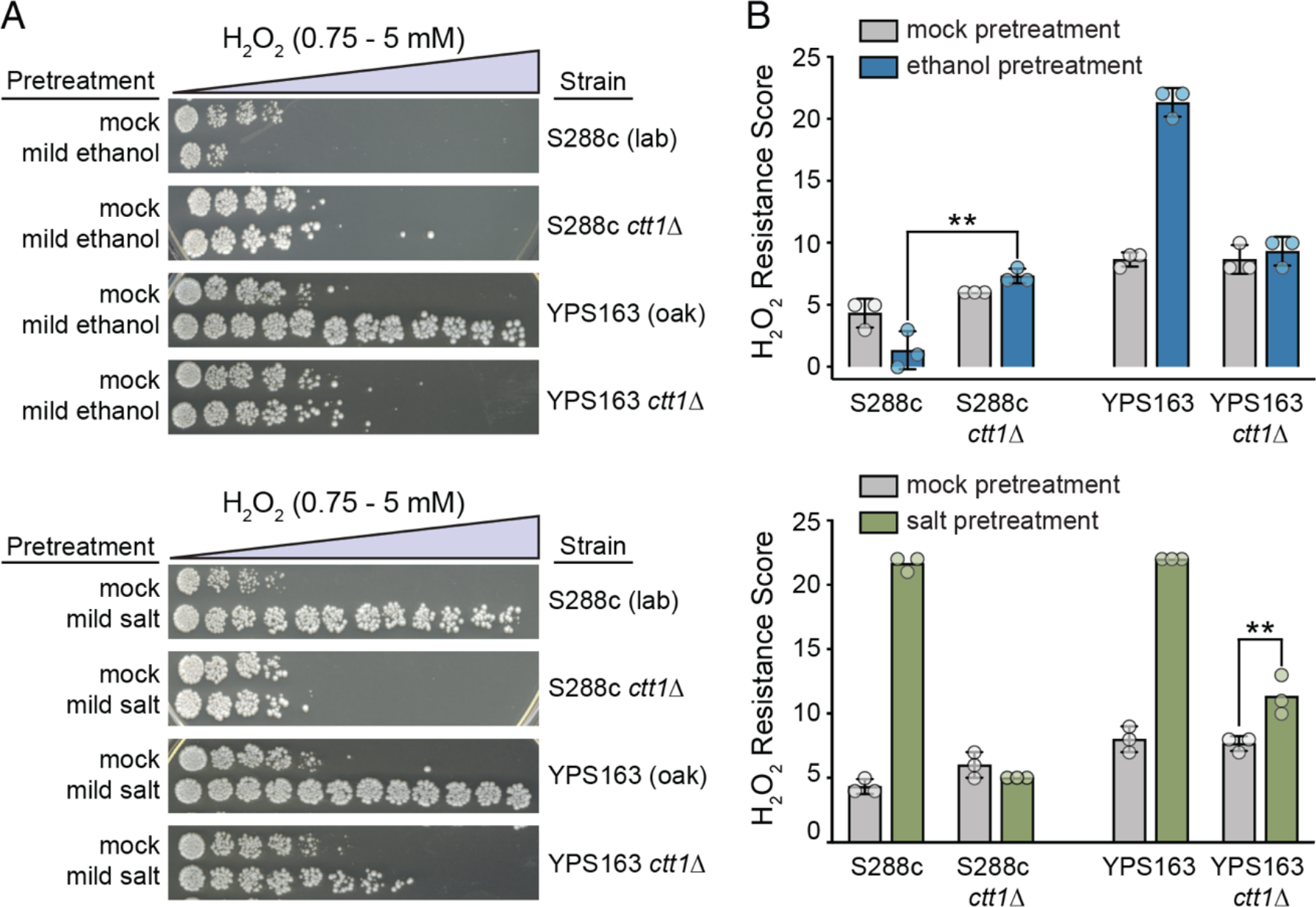
CTT1 is partially dispensable for salt-induced, but not ethanol-induced, cross protection against H2O2 in a wild oak strain. (A) Representative acquired H2O2 resistance assays for wild-type S288c (lab) and YPS163 (oak) strains and respective *ctt1*Δ mutants. Strains were exposed to a mild ‘primary’ stress pretreatment (5% ethanol or 0.4 M NaCl in culture media) or a mock control for 60 minutes, collected and resuspended in fresh media, exposed to a panel of 11 doses of severe ‘secondary’ doses of H2O2 for 2 hours, and then spot plated to score viability. (B) A single survival score was calculated from the viability at all H2O2 doses (see Materials and Methods). Each plot shows the mean and standard deviation of three independent biological replicates. The replicates for some strain conditions all had the same tolerance score and thus zero standard deviation (see S3 Table for raw numerical data). Acquired H2O2 resistance was significantly higher in the YPS163 *ctt1*Δ mutant with NaCl as the pretreatment compared to the mock control (** *P <* 0.01, ordinal regression analysis on raw spot scores—see Methods), while the sensitizing effect of ethanol-pretreatment on S288c was eliminated in the S288c *ctt1*Δ mutant (** *P* < 0.01, ordinal regression analysis on raw spot scores).

To better understand the scope of *CTT1* conditional dependency, we deleted *CTT1* in a panel of yeast strains from diverse ecological niches (Table S1), and then assayed acquired H2O2 resistance following either mild ethanol or salt pretreatments. With the exception of a wild coconut strain (Y10), lack of *CTT1* had little to no effect on intrinsic (basal) levels of H2O2 tolerance in the absence of pretreatment (Figure 2A and Figure S1). For cross protection however, we found that *CTT1* was partially dispensable in approximately half of tested strains (6/13), with half of those displaying *CTT1-*independent cross protection in only one condition (always salt-induced cross protection) (Figure 2B). Hierarchical clustering of the tolerance scores showed that oak and vineyard strains partitioned together (Figure 2C), potentially providing another example of trait differentiation between oak and vineyard populations [55], though larger sample sizes are certainly needed to determine whether this observation is generalizable. To understand whether the lack of ethanol-induced cross protection against H2O2 was correlated with *CTT1* dependency for salt-induced cross protection, we included two strains (YJM1129 and YJM627) that, similar to S288c, displayed low levels of acquired H2O2 resistance following mild ethanol-pretreatment [50]. *CTT1* was partially dispensable for salt-induced cross protection in one of the two strains (Figure 2B), suggesting that the absence of ethanol-induced cross protection was uncorrelated with *CTT1* dispensability when salt was the primary stress. In summary, *CTT1* dispensability demonstrates complex patterns of GxE interactions across genetically diverse wild yeast strains, while suggesting the existence of multiple molecular pathways and/or mechanisms that can lead to superficially similar phenotypes (in this case maximal acquired stress resistance).

**Figure 2.**
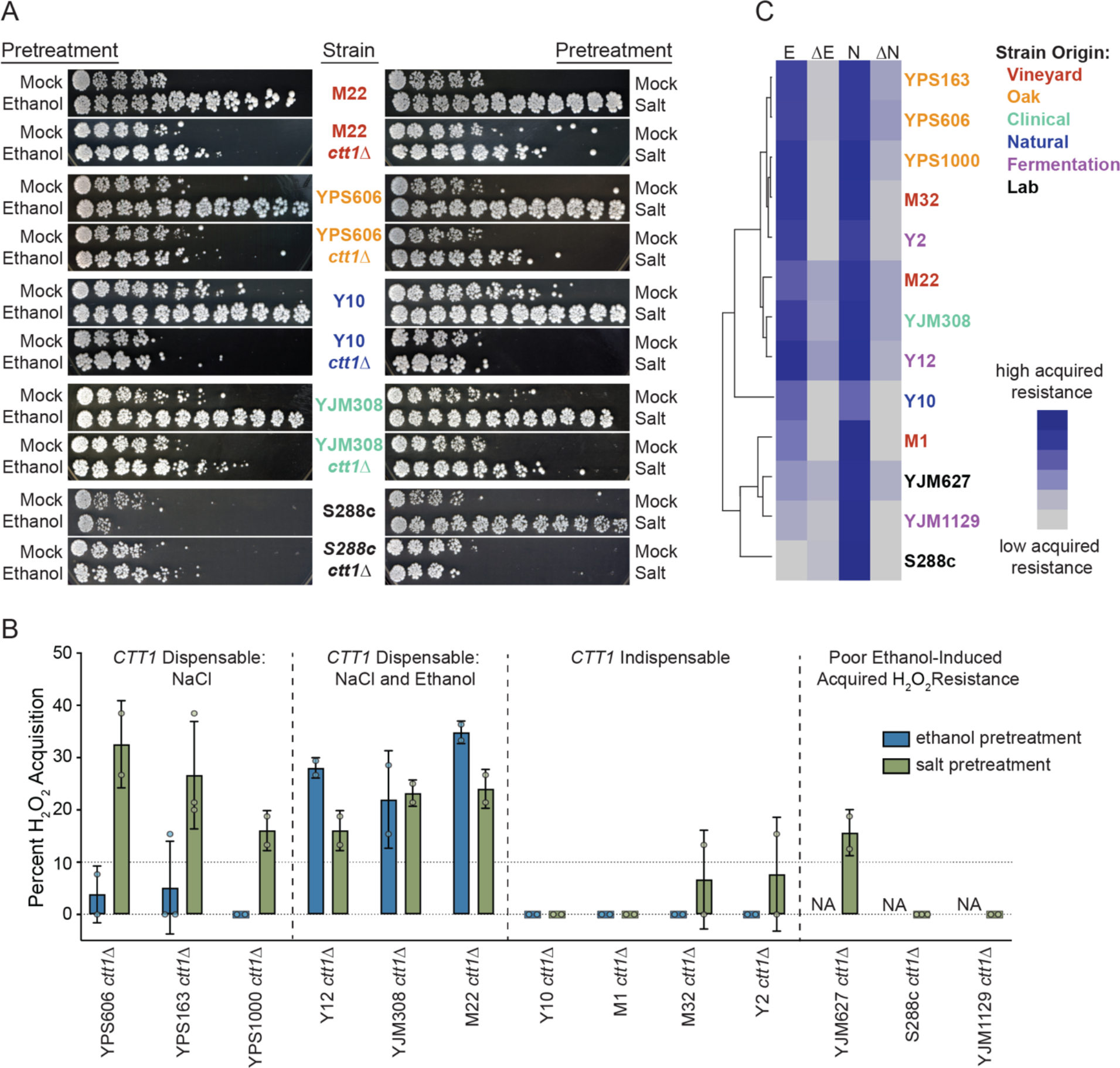
Diverse yeast strains show complex patterns of conditional *CTT1* dependency. (A) Representative acquired H2O2 resistance assays of wild-type strains and their *ctt1*Δ mutant derivatives. (B) Percent acquired H2O2 resistance in *ctt1Δ* mutants relative to that of the wild type revealed three classes of conditional *CTT1* dependency (based on a cutoff of 10% or higher residual acquisition, see Methods). A fourth class represents strains with poor acquired H2O2 resistance following ethanol pretreatment. Error bars denote the mean and standard deviation of biological duplicates, except for S288c and YPS163 which are in biological triplicate. (C) Hierarchical clustering of H2O2 survival scores for wild-type strains following either ethanol (E) or salt (N) pretreatments, or their *ctt1Δ* derivatives (ΔE and ΔN). Strain labels are color coded according to their origin as indicated in the key on the right.

### Extensive variation in the ethanol and salt responses across yeast strains

Stress-activated gene expression changes are necessary for acquired stress resistance [45, 48], and natural variation in cross protection has been linked to gene expression variation [50]. Therefore, we performed transcriptional profiling in a subset of strains with different levels of *CTT1* dispensability to identify differentially expressed genes and their potential transcriptional regulators that may be important for the phenotypic differences in cross protection. We included two each of strains with poor ethanol-induced acquisition (S288c and YJM1129), *CTT1* indispensability for both NaCl and ethanol pretreatments (M1 and Y10), *CTT1* dispensability for only NaCl pretreatment (YPS163 and YPS606), and *CTT1* dispensability for both NaCl and ethanol (M22 and YJM308). RNA-seq was performed on these strains comparing an unstressed control to cells treated with either 5% v/v ethanol for 30 minutes or 0.4 M NaCl for 45 minutes (timepoints which encompass the peak response for each stressor [45, 48]).

Differential expression analysis was performed by comparing the log2 fold changes during stress for each strain to the mean expression for all 8 strains. Out of the 6,092 yeast genes expressed in at least one condition (cpm ≥ 1), 4,261 genes (69%) showed significant differences in ethanol-responsive expression in at least one strain compared to the mean of all strains, while 3,197 (54%) were differentially expressed in at least one strain during the NaCl response. To identify modules of co-regulated genes that differ in expression across strains, we performed hierarchical clustering on both strains and significantly differentially-expressed genes (see Methods). For ethanol stress, strains with poor overall cross protection (S288c and YJM1129) were clear outliers. S288c in particular had lower induction of stress defense-enriched clusters (Figure 3A, rightmost column). While YJM1129 was also an outlier compared to other strains, lower overall induction of stress defense genes was not generally observed.

**Figure 3.**
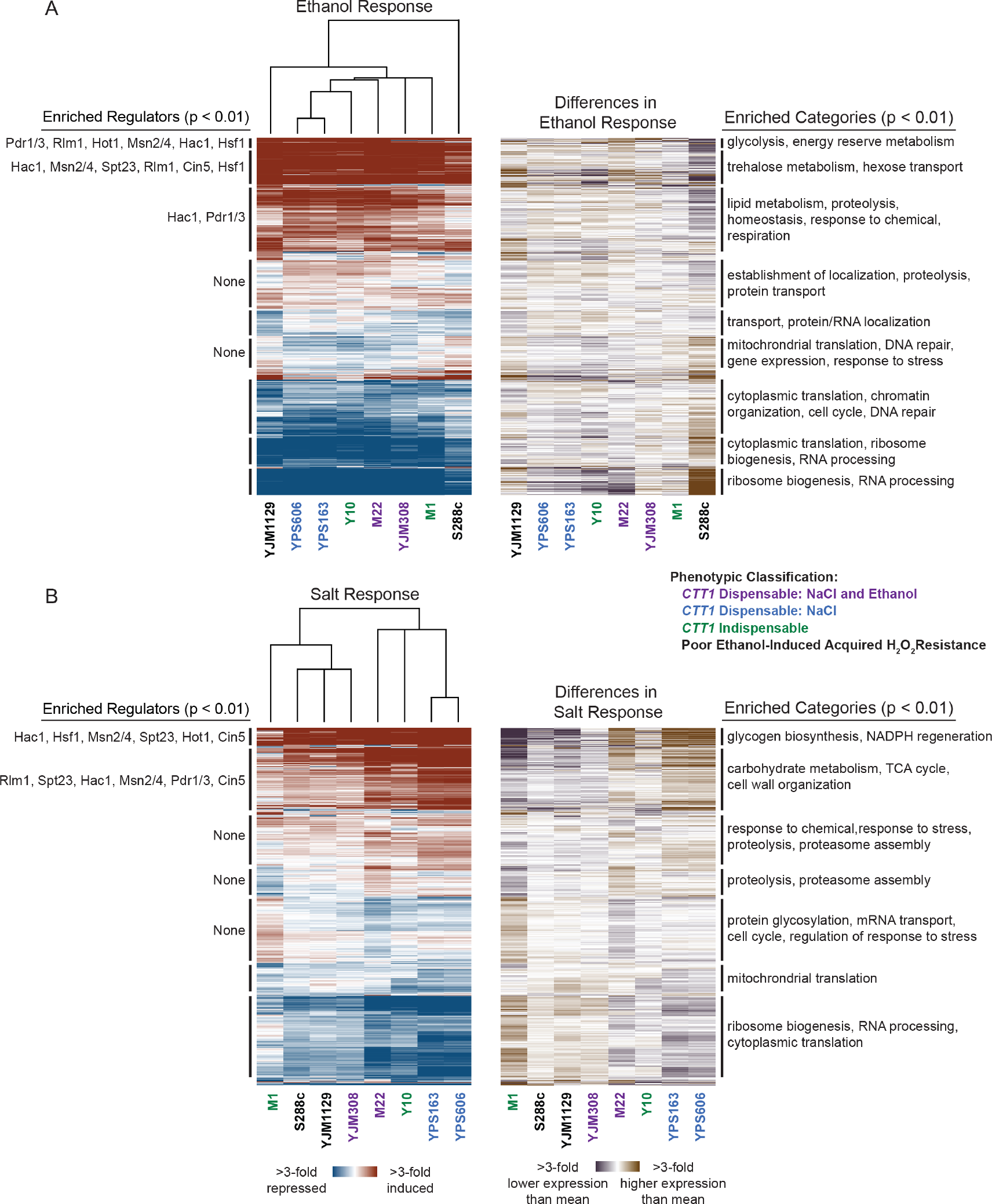
**Extensive variation in stress responsive gene expression across diverse yeast isolates**. Hierarchical clustering of 4,262 genes with significant (FDR < 0.01) differential ethanol-responsive expression in at least one strain relative to the mean of all strains (A), and 3,918 genes with significant (FDR < 0.01) differential salt-responsive expression in at least one strain relative to the mean of all strains (B). Genes are shown as rows and log2 fold changes for each strain’s stress response (left) or difference in stress response (right) are shown as columns. Red indicates induced and blue indicates repressed expression in response to stress, while violet indicates higher and brown indicates lower expression relative to the mean stress response of all strains. Significant regulatory enrichments are annotated to the left and functional enrichments are annotated to the right (Bonferroni-corrected *P* < 0.01). Full lists of regulatory and functional enrichments are in Table S7.

Moreover, while the lack of ethanol-induced cross protection in S288c is likely due to reduced *CTT1* mRNA expression (12-fold lower relative abundance during ethanol stress compared to the mean of all strains), this is likely not true of YJM1129 (which had only 1.5-fold lower relative mRNA abundance during ethanol stress compared to the mean of all strains), suggesting a different mechanistic basis for the absence of ethanol-induced cross protection in this strain (see Discussion).

Strains did not generally cluster together based on their *CTT1* dispensability phenotype, suggesting a weak overall correlation between stress-responsive gene expression and these specific acquired resistance phenotypes. This is perhaps not surprising, considering the complexity of genomic responses to stress and their likely effects on multiple traits [56, 57].

However, strain-specific differences in stress-responsive expression were clearly apparent. For example, YPS163, YPS606, Y10, and M22 all had higher expression for salt-induced clusters that were functionally enriched for stress related processes such as in proteolysis (*P* = 4 x 10^-9^), cell wall organization (*P* = 2 x 10^-5^), and response to stress (*P* = 2 x 10^-4^) (Figure 3B). Likewise, M1 had decreased expression for those same clusters, while having novel or amplified induction during salt stress for a cluster functionally enriched for genes involved in the cell cycle (*P* = 5 x 10^-11^), protein glycosylation (*P* = 2 x 10^-5^), mRNA transport (*P* = 8 x 10^-4^), and regulation of the response to stress (*P* = 1 x 10^-3^).

To explore strain-specific differences in stress responsive gene expression further, we identified genes in each strain that differed from the mean response of all strains (i.e., the ‘consensus’ response, see Methods), and separated them into classes (e.g., novel induction, amplified repression, Figure 4). As predicted from the clustering and consistent with our previous studies comparing S288c to a smaller panel of strains [48, 58], S288c had the largest number of genes with divergent expression compared to the consensus for ethanol stress (∼600 genes with defective induction and ∼700 genes with defective repression). These aberrantly expressed genes were functionally enriched for several categories related to stress defense (Table 1). Interestingly, YJM1129, which does not acquire further H2O2 resistance following ethanol pretreatment, did not have a particularly high number of genes (107) with lower induction during ethanol stress, though there was a small enrichment for response to environmental stimulus (*P* = 2 x 10^-4^). M1, despite not having a discernable defect in acquired stress resistance under the conditions tested, had a large number (∼400) of salt-induced genes with lower induction than consensus, and these were enriched for carbohydrate metabolism (*P* = 2 x 10^-17^), trehalose metabolism (*P* = 7 x 10^-9^), and response to chemical (*P* = 9 x 10^-3^).

**Figure 4.**
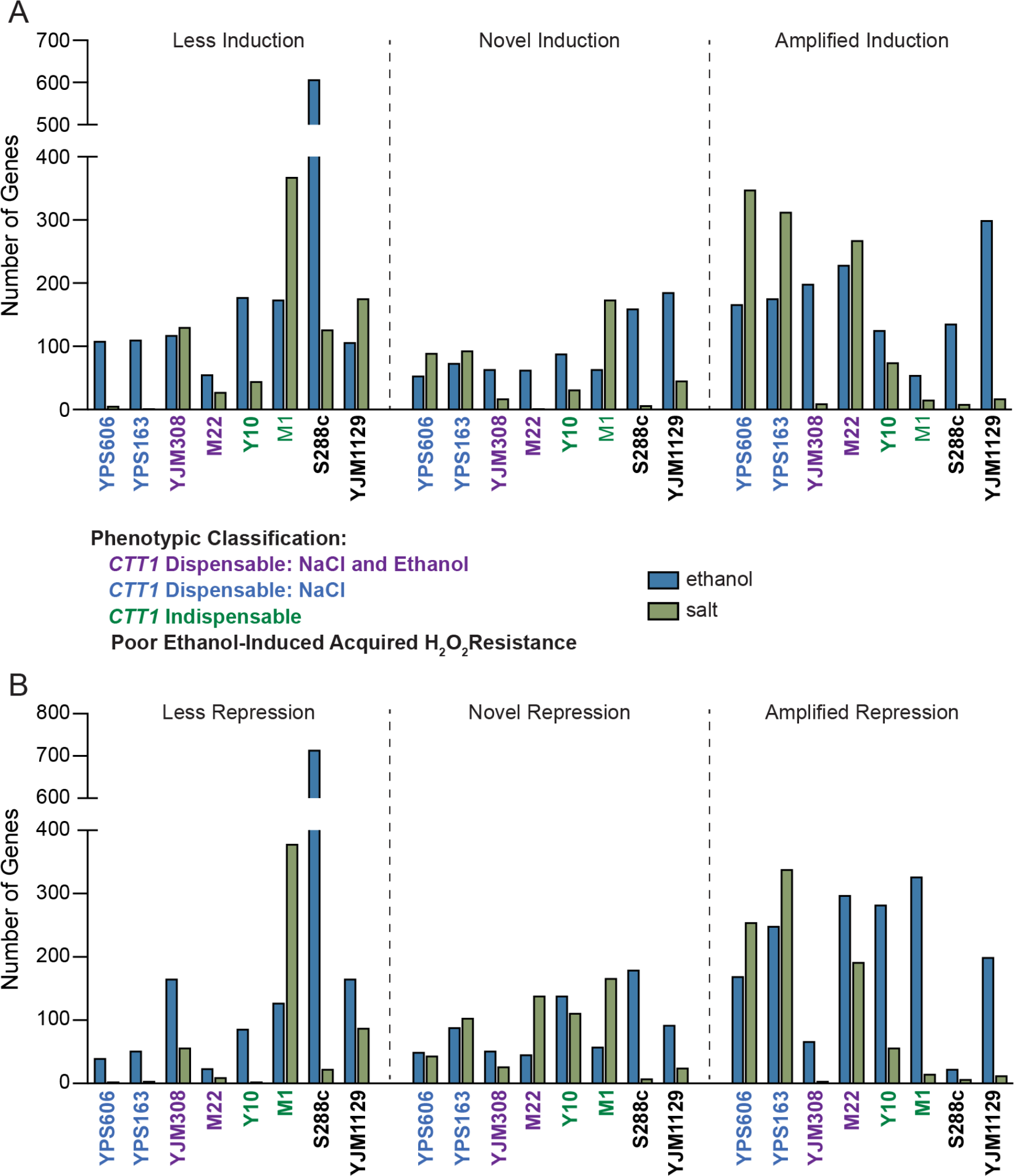
Strain-specific variation in different classes of stress-activated expression changes. Total number of genes for each strain with significantly (FDR < 0.01) different (> 1.5- fold) induction patterns (A) or repression patterns (B) relative to the mean or ‘consensus’ response (see Methods). Tables S8 and S9 contain the genes for ethanol-responsive and salt- responsive categories, respectively.

**Table 1.**
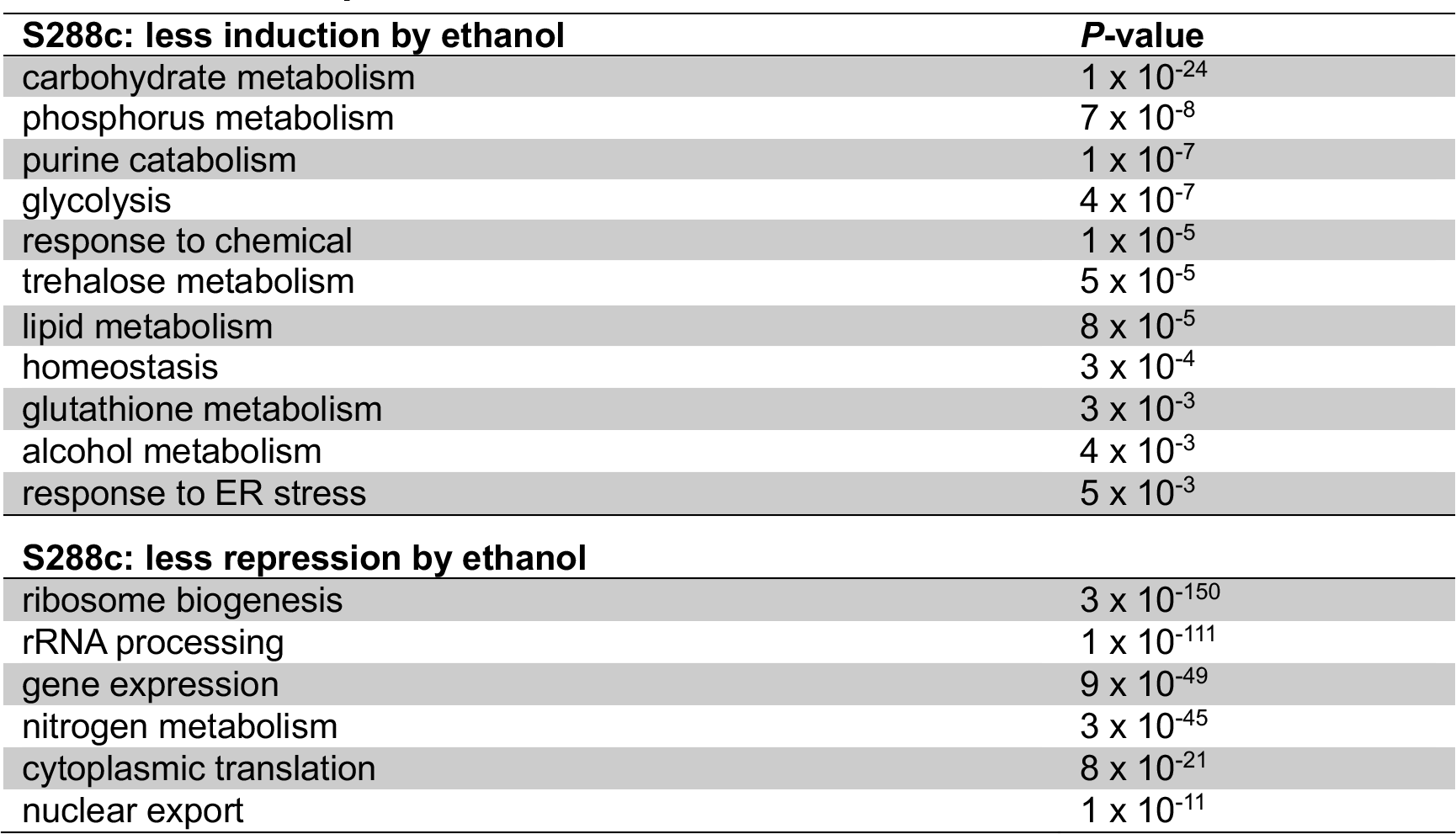
Functional enrichments of genes where the S288c ethanol response differs from the consensus response.

We were particularly interested in strains with amplified stress-induced expression compared to all strains, as this may indicate a heightened response. Surprisingly, despite being deficient in ethanol-induced cross protection against H2O2, YJM1129 had the most genes (300) with amplified induction during ethanol stress, though most were not functional enriched for processes related to stress defense (Table 2). For salt stress, the oak strains had the highest number of genes with amplified induction (313 genes for YPS163 and 348 for YPS606), with the vast majority of those genes being shared between the two strains (258), which is unsurprising considering the similarity of their expression responses. Shared genes with amplified induction in YPS163 and YPS606 during salt stress were enriched for functions implicated in stress defense (Table 2), leading us to hypothesize that a subset of those genes with amplified induction are responsible for the residual salt-induced acquired H2O2 resistance in the absence of *CTT1* in the oak strain background (see below).

**Table 2.** Functional enrichments of genes with amplified induction during stress relative to the consensus response.

| **YJM1129: amplified induction by ethanol** |  |
| --- | --- |
| small molecule metabolism | $1 \times 10^{-8}$ |
| cell wall organization | $3 \times 10^{-4}$ |
| respiration | $4 \times 10^{-3}$ |
| carbohydrate metabolism | $5 \times 10^{-3}$ |
| fatty acid catabolism | $6 \times 10^{-3}$ |
| **YPS163 and YPS606: amplified induction by salt** |  |
| carbohydrate metabolism | $1 \times 10^{-16}$ |
| trehalose metabolism | $1 \times 10^{-8}$ |
| pyrimidine metabolism | $4 \times 10^{-7}$ |
| phosphorus metabolism | $3 \times 10^{-4}$ |
| response to chemical | $2 \times 10^{-3}$ |
| arginine biosynthesis | $4 \times 10^{-3}$ |
| glycogen biosynthesis | $8 \times 10^{-3}$ |

### Identification of regulators necessary for ethanol or salt induced cross protection against H2O2

To identify potential regulators of the salt and ethanol responses that may be responsible for acquired hydrogen peroxide resistance, we looked for regulatory enrichments for genes belonging to stress-induced clusters (Figure 3 and Table S7). Prioritizing based on regulatory enrichment as well as transcription factors with known functions in relevant stressors (ethanol, osmotic, and/or oxidative stresses), we generated eight transcription factor gene deletions in the YPS606 strain background (*hot1Δ*, *HSF1*/*hsf1Δ*, *msn1Δ*, *msn2Δmsn4Δ, skn7Δ*, *sko1Δ*, *smp1Δ*, *yap1Δ*; see Table 3 for a brief description of each transcription factor), and tested whether those regulators were necessary for cross protection. We also tested the transcription factor mutations in combination with *ctt1*Δ mutations, to determine whether the tested transcription factors were potential regulators of the putative alternative peroxidases necessary for *CTT1*-independent acquisition. We chose YPS606 as a representative strain for further genetic dissection for two reasons. First, YPS606 possessed the highest level of residual acquisition in the absence of *CTT1*, providing the largest dynamic range for detecting loss of *CTT1*-independent acquisition. Second, we reasoned that because *CTT1* is only partially dispensable for salt-induced cross protection and not for ethanol, comparison of expression profiles for potential compensatory peroxidases during both ethanol and salt stress may help implicate the responsible party.

**Table 3.** Potential transcription factors involved in acquired stress resistance.

| **Transcription Factor (TF)** | **Description** | **References** |
| --- | --- | --- |
| Hot1p | Osmotic stress responsive TF | [84] |
| Hsf1p | Essential heat shock factor | [85, 86] |
| Msn1p | Osmotic stress responsive TF | [84] |
| Msn2/4p | Paralogous general stress-responsive TFs | [86-88] |
| Skn7p | Osmotic and oxidative stress-responsive TF | [89-91] |
| Sko1p | Osmotic stress-responsive TF | [92] |
| Smp1p | Osmotic stress responsive TF | [93] |
| Yap1p | Oxidative stress-responsive TF | [91] |

With ethanol as the pretreatment, the mutant with the largest defect in acquired resistance was the *msn2Δ msn4Δ* double mutant (Figure 5A), where cross protection was completely abolished. The *skn7*Δ mutant had an intermediate defect, as well as a slight defect for intrinsic (basal) H2O2 resistance. The *yap1Δ* mutant had a strong intrinsic H2O2 resistance defect, but was still able to fully acquire resistance, suggesting that low intrinsic stress resistance does not necessarily preclude maximal acquisition. In contrast to the *yap1Δ* and *skn7Δ* single mutants, the *yap1Δ ctt1Δ* and *skn7*Δ *ctt1Δ* double mutants were unable to acquire further H2O2 resistance beyond their lowered intrinsic H2O2 resistance (Figure 5A), suggesting that Yap1p and Skn7p may regulate alternative peroxidases responsible for residual acquired H2O2 resistance in the absence of Ctt1p (see below).

**Figure 5.**
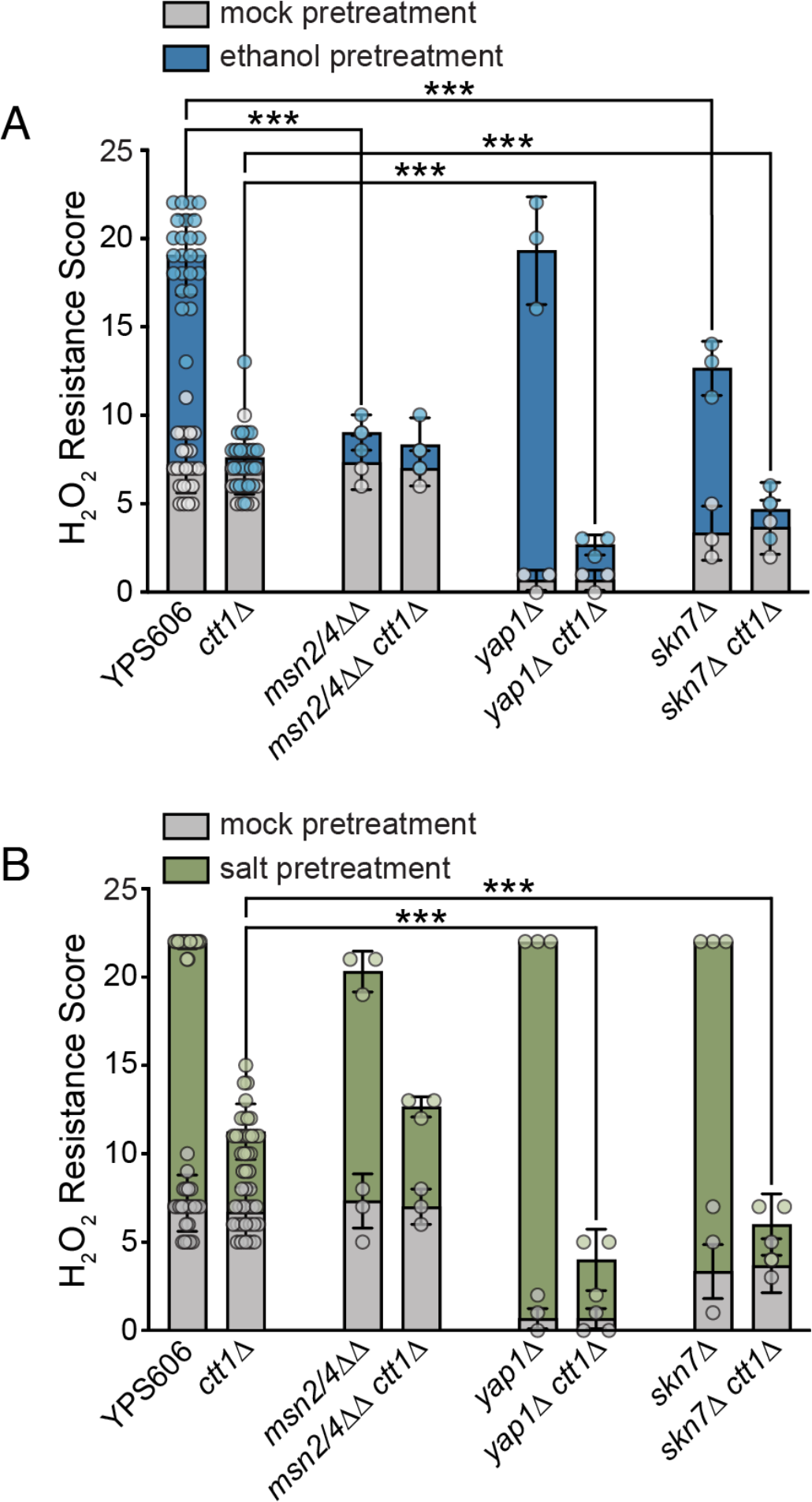
*msn2Δ msn4Δ*, *skn7*Δ, and *yap1Δ* mutants have varied effects on intrinsic and acquired H2O2 resistance. Survival score plots representing the mean and standard deviation for biological triplicates for transcription factor mutant strains with either 5% ethanol (A) or 0.4 M NaCl (B) as the mild pretreatment. Because each transcription factor mutant was tested alongside a wild-type YPS606 and YPS606 *ctt1Δ* control including those shown in Figure S2, those bar graphs depict 24 replicates across all experiments. The replicates for NaCl-treated *yap1*Δ and *skn7Δ* mutants all had the same tolerance score and thus zero standard deviation (see Tables S3 and S4 for raw numerical data). Asterisks represent significant differences in acquired resistance between denoted strains (*** *P* < 0.001 ordinal regression analysis on raw spot scores).

For salt as the mild pretreatment, none of the tested mutants showed a defect in acquired H2O2 resistance when *CTT1* was present (Figure 5B and Figure S2), which was especially striking considering that *msn2Δ msn4Δ* double mutants have extremely low salt-induced cross protection against H2O2 in the S288c background [41, 45]. This is likely due to partial regulatory redundancy in the YPS606 strain background (see below). Intriguingly, both the *yap1*Δ *ctt1Δ* and *skn7*Δ *ctt1Δ* double mutants had significantly reduced levels of *CTT1*- independent acquisition (Figure 5B), suggesting that both Yap1p and Skn7p play a role in regulating those compensatory functions. Because low intrinsic stress resistance does not preclude the ability to acquire high levels of stress resistance, it was somewhat striking that both the *skn7*Δ *ctt1Δ* and especially the *yap1*Δ *ctt1Δ* had little acquisition beyond that of the intrinsic resistance of the single *skn7*Δ or *yap1Δ* mutants. This suggests that the peroxidases necessary for intrinsic resistance may overlap with those that partially compensate in the absence of Ctt1p. None of the other transcription factors tested (*HOT1*, *HSF1*, *MSN1*, *SKO1, or SMP1*) showed a defect in acquired resistance under any condition (Figures S2 and S3), though *HSF1* comes with the caveat that we were only able to test a heterozygous deletion due to *HSF1* being essential.

### Transcriptional profiling reveals the overlapping and unique roles of the Msn2/4p, Skn7p, and Yap1p transcription factors during either ethanol or salt stress

We next performed expression profiling on transcription factor mutants in the YPS606 background that had an acquired stress resistance defect (either in isolation or in combination with *ctt1*Δ) for both mild ethanol and mild salt stresses, as well as for an unstressed control: *msn2Δ msn4Δ*, *skn7Δ*, and *yap1Δ*. For the ethanol response, the *msn2Δ msn4Δ* mutant had the largest number of genes with significantly defective induction (i.e., reduced induction relative to wild type)—379 genes. In contrast, 95 genes showed defective induction in the *skn7Δ* mutant, while only 28 genes showed defective induction in the *yap1Δ* mutant (Figure 6 and Table S6). We next characterized the extent of overlap for the induced ethanol response regulons for each transcription factor (Figure 6). Eight genes were part of the ethanol-induced regulon for all 3 transcription factors, including four genes involved in protein folding (*AHA1, HSP78, SIS1, ERO1;* enrichment *P* = 4 x 10^-4^), as well as genes encoding an alcohol dehydrogenase (*ADH5*), a meiotic regulator *(WTM1*), a hexokinase (*EMI2*), and stress-responsive protein of unknown function (*HOR7*). There were 22 genes in the overlapping *SKN7-YAP1* regulons, which were enriched for amino acid metabolism (*P* = 1 x 10^-3^). There was no significantly enriched functional category for the genes that were part of the overlapping *SKN7* and *MSN2/4* regulons, though *CTT1* was one of those genes. In fact, ethanol-responsive fold-changes in *CTT1* expression correlated well with the total (*msn2Δ msn4Δ*) and partial (*skn7Δ*) cross protection defects in those mutants—*CTT1* showed no induction (1.3-fold) in the *msn2Δ msn4Δ* mutant and 32-fold induction in the *skn7Δ* mutant (compared to 91-fold induction in the wild-type strain). In contrast, *CTT1* was 136-fold induced in the *yap1Δ* strain, which had wild-type levels of acquired resistance, suggesting that Yap1p plays no role in *CTT1* regulation during ethanol stress. Finally, there were 325 genes that were unique to the *MSN2/4* ethanol regulon, which were enriched for stress related functions such as metabolism of trehalose (*P* = 6 x 10^-6^) and glycogen (*P* = 3 x 10^-4^), and the response to oxidative stress (*P* = 5 x 10^-3^).

**Figure 6.**
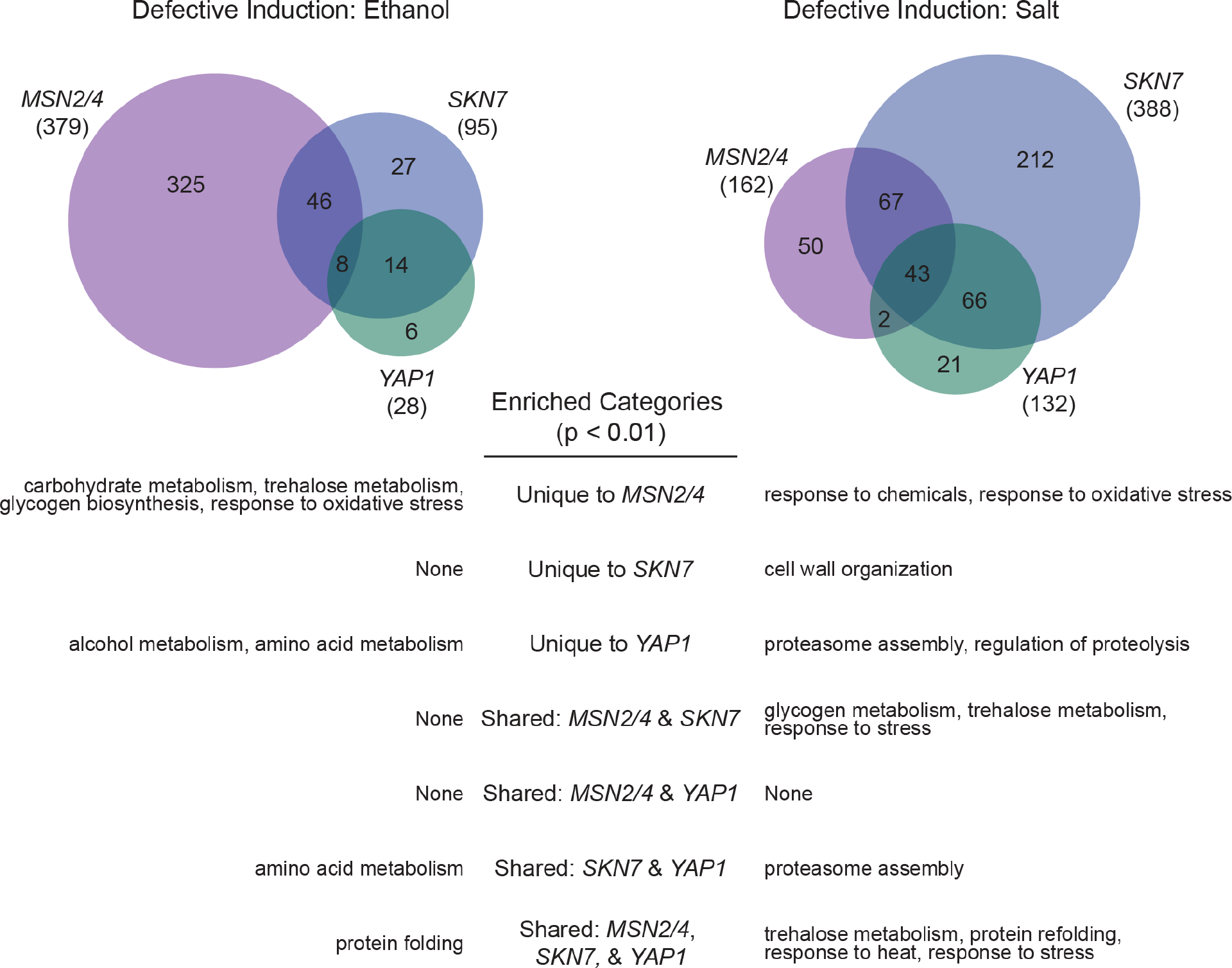
Msn2/4, Skn7, and Yap1 regulate shared and unique targets for the YPS606 ethanol and NaCl responses. The Venn diagrams represent genes with significant defective induction for each transcription factor mutant’s (*msn2Δmsn4Δ, skn7Δ*, or *yap1Δ*) stress response (FDR < 0.01 and at least 1.5-fold reduced expression). Significant functional enrichments (Bonferroni-corrected *P* < 0.01) are shown below for all unique and shared sets of regulated genes.

In contrast with the ethanol response, *SKN7* had the largest salt-induced regulon (388 genes), while *MSN2/4* and *YAP1* had moderately sized regulons (162 and 132 genes, respectively). There were 43 genes with that were part of the salt-induced regulons for all three transcription factors, which were enriched for response to stress (*P* = 5 x 10^-3^), protein refolding (*P* = 1 x 10^-9^), and trehalose metabolism (*P* = 9 x 10^-9^). *CTT1* was part of this overlapping group, though the *CTT1* is much more strongly induced by salt than ethanol in the wild-type strain (340-fold vs. 91-fold) and still possessed relatively high induction in all three mutants (63-fold for *msn2Δ msn4Δ*, 94-fold for *skn7Δ*, and 63-fold for *yap1Δ*). *CTT1* expression thus appears combinatorially controlled by Msn2/4p, Skn7p, and Yap1p, which would explain why cells were still able to acquire resistance in the absence of any single transcription factor.

While the overlap in regulons was more substantial for salt, each transcription factor also regulated at least 20 genes uniquely. The 212 genes that were unique to the *SKN7* salt-induced regulon were enriched for cell wall organization (*P* = 8 x 10^-8^), the 50 genes unique to the *MSN2/4* salt-induced regulon were enriched for responses to chemical (*P* = 1 x 10^-3^) and oxidative (*P* = 2 x 10^-3^) stresses, and the 21 genes unique to the *YAP1* salt regulon were enriched for proteasome assembly (*P* = 2 x 10^-5^) and regulation of proteolysis (*P* = 4 x 10^-4^).

There were 110 genes in the overlapping *SKN7-MSN2* regulons, which were enriched for the response to stress (*P* = 3 x 10^-3^), and glycogen (*P* = 6 x 10^-7^) and trehalose (*P* = 1 x 10^-14^) metabolism. There were 22 genes in the overlapping *SKN7-YAP1* regulons, which were enriched for proteasome assembly (*P* = 3 x 10^-4^). Finally, there were no shared functionally enriched categories for the shared *MSN2-YAP1* regulons.

For intrinsic (basal) gene expression, there were 197 genes with differential expression compared to wild type in the *yap1Δ* mutant, 369 genes in the *skn7Δ* mutant, and 10 genes in the *msn2Δ msn4Δ* mutant (Table S6). For the *yap1Δ* strain, 64 genes had at least 1.5-fold higher expression relative to the wild-type strain (in other words, were de-repressed), which were enriched for amino acid biosynthesis (*P* = 7 x 10^-12^). Fifty-six genes had >1.5-fold lower expression in the *yap1Δ* strain (i.e., defective activation), which were importantly enriched for cellular oxidant detoxification (*P* = 3 x 10^-8^). This includes genes that encode for proteins known to be involved in directly detoxifying H2O2 (*GPX2, PRX1, GRX2, AHP1, TSA1*) or genes necessary for proper function of those systems (*GSH1* involved in glutathione biosynthesis and *TRR1* encoding a thioredoxin reductase). Decreased expression of these genes likely explains the strong intrinsic H2O2 tolerance defect of the *yap1*Δ strain. The *skn7Δ* mutant, which also had an intrinsic H2O2 tolerance defect, had 44 genes with >1.5-fold lower expression than wild type. Those 44 genes were also enriched for cellular oxidant detoxification (*P* = 4 x 10^-4^), including genes encoding direct H2O2 detoxifiers (*PRX1, GPX2, TSA1, AHP1*). There was strong and significant overlap between genes with either lower expression in both the *yap1Δ* and *skn7*Δ strains (44-fold over-enrichment and *P* = x 10^-26^, Fisher’s exact test) or higher expression (31- fold over-enrichment and *P* = 6 x 10^-77^, Fisher’s exact test), suggesting substantial overlap for the two regulons. Genes with >1.5-fold higher expression in both the *yap1*Δ and *skn7Δ* mutants relative to wild type were enriched for amino acid biosynthesis (*P* = 2 x 10^-13^). The 99 genes with higher expression in the *skn7Δ* mutant but not the *yap1Δ* mutant were enriched for those involved in iron ion transport (*P* = 3 x 10^-5^).

In summary, Msn2/4p play the largest role amongst the transcription factors tested for regulating the YPS606 ethanol response, while Skn7p plays the largest role in regulating the YPS606 salt response. During ethanol stress, Msn2/4p uniquely regulate genes enriched for key stress defense processes, which correlates with the absence of ethanol-induced acquired H2O2 resistance in the *msn2Δ msn4Δ* mutant. In contrast, there was far more overlap between the Skn7p, Msn2/4p, and Yap1p regulons for salt stress—including coordinate regulation of key stress defense processes—which correlates with the observation wild-type levels of salt-induced acquired H2O2 in mutants lacking a single regulator.

### *CTT1*-independent acquisition requires glutathione

Finally, we sought to understand the mechanistic basis for partial *CTT1* dispensability. Leveraging our transcriptomic data, we focused on the expression of genes annotated to have any kind of peroxidase activity. We clustered just the expression of those genes (Figure 7), which revealed interesting patterns of differential expression across strains where *CTT1* was either dispensable or indispensable for acquisition. Clustering on strains for just this subset of genes better matched acquired resistance phenotypes, particularly with ethanol, where S288c clearly showed reduced expression of most peroxidases in addition to *CTT1*, while the strains where *CTT1* was partially dispensable for ethanol (M22 and YJM308) clustered together and showed higher expression for most peroxidase genes. For salt, strains where *CTT1* was indispensable for acquisition (M1 and Y10, plus S288c and YJM1129) tended to have lower than average induction of peroxidase genes, with S288c and YJM1129 in particular clustering together.

**Figure 7.**
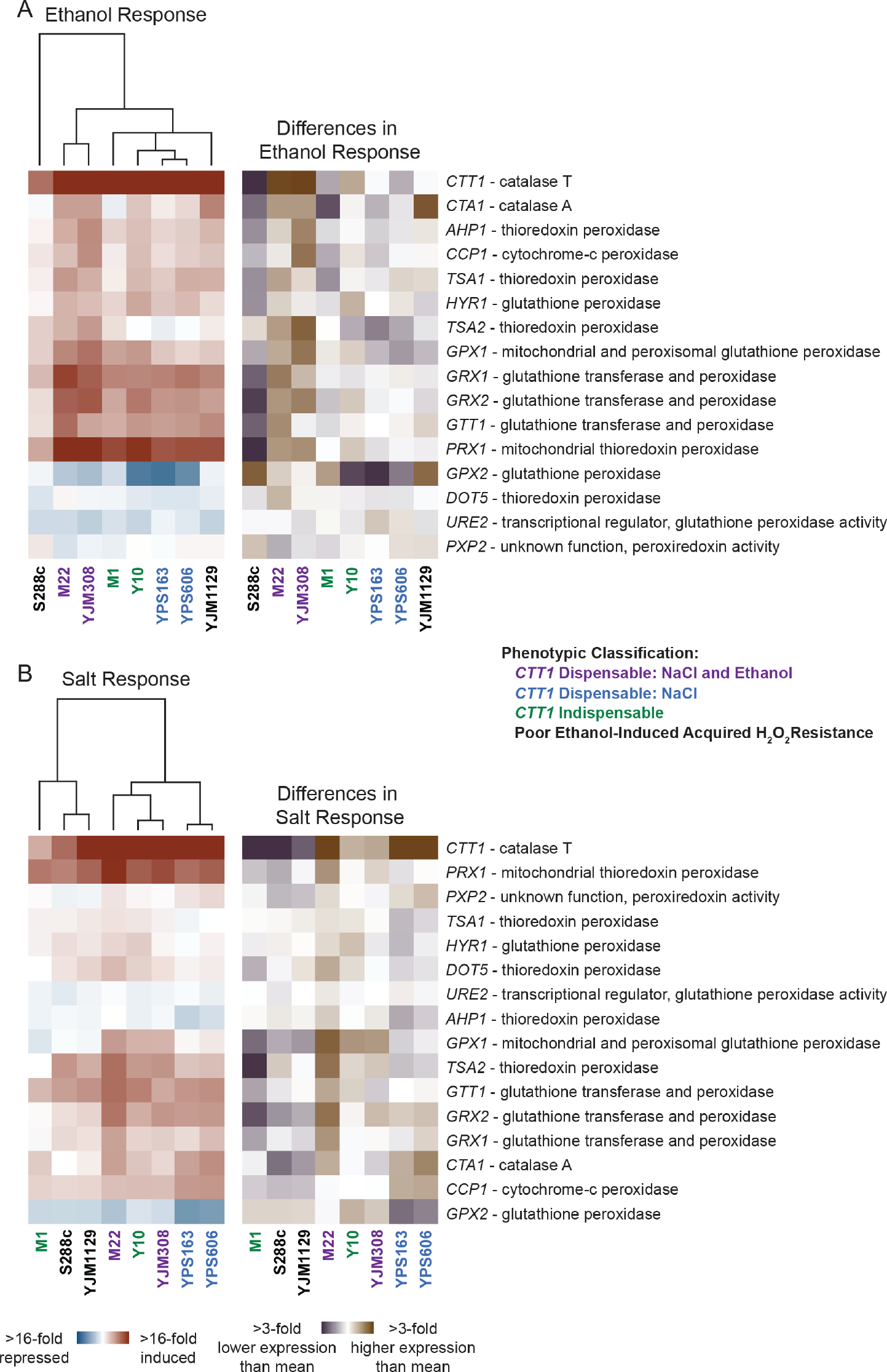
Stress-responsive expression variation in genes involved in detoxifying reactive oxygen species. Hierarchical clustering of all yeast genes annotated as having peroxidase activity in response to either ethanol (A) or NaCl (B). Genes are shown as rows and log2 fold changes for each strain’s stress response (left) or difference in stress response (right) are shown as columns. Red indicates induced and blue indicates repressed expression in response to stress, while violet indicates higher and brown indicates lower expression relative to the mean stress response of all strains.

For all non-*CTT1* peroxidase genes that were significantly induced at least 2-fold in any strain, expression was higher in strains where *CTT1* was dispensable (1.9-fold higher induction for ethanol, *P* < 4 x 10^-4^, t-test, and 1.6-fold higher for salt, *P* < 4 x 10^-3^, t-test), suggesting potentially redundant mechanisms. Because there were multiple, potentially redundant candidate genes that could be responsible for *CTT1*-independent acquisition, we sought to narrow the possible mechanisms. In addition to catalases, the major cellular peroxidases require reduction by either thioredoxin or glutathione to continue scavenging H2O2. Intriguingly, genes encoding glutathione-dependent peroxidases (*GRX2, GPX1, GPX2*) were among the those with the largest differences in expression between *CTT1-*dispensable and *CTT1-* indispensable strains during either ethanol or salt stresses. Additionally, the YPS606 *skn7Δ* mutant, which lacks *CTT1*-indepdent acquisition, also showed defective induction during salt stress for several glutathione-dependent peroxidase genes (*GPX1, GRX2, GRX1*). And while genes encoding glutathione-dependent peroxidases did not have lower expression in the *yap1*Δ mutant during salt stress, we did notice that expression of the gene that encodes the first enzymatic step of glutathione biosynthesis (*GSH1*) was 1.9-fold lower, suggesting that the *yap1Δ* mutant may have insufficient glutathione for those peroxidases to properly function.

Altogether, these data suggested that the peroxidase(s) responsible for *CTT1*- independent acquisition may be glutathione dependent. Thus, we tested the effects of loss of *GSH1* on acquired stress resistance, both alone and in combination with *ctt1Δ* in one strain that possesses *CTT1*-independent acquisition following either ethanol or salt pretreatments (M22) or two strains that only possess *CTT1*- independent acquisition following salt pretreatment (YPS163 and YPS606). Each *gsh1*Δ single mutant was able to fully acquire hydrogen peroxide resistance for either ethanol or salt pretreatments (Figure 8), suggesting that *GSH1* is not necessary for acquisition in the presence of *CTT1*. However, lack of *GSH1* significantly reduced *CTT1*-independent acquisition for M22 under both stress pretreatments (Figure 8), as well as for YPS163 and YPS606 during salt pretreatment (Figure 8B). For M22, the *gsh1Δ ctt1Δ* mutant also had an intrinsic H2O2 resistance defect in addition to the loss of *CTT1*- independent acquisition, suggesting that the compensatory peroxidases and *CTT1* may play a role in both processes. Overall, the data suggest that glutathione-dependent peroxidases are at least partially responsible for the conditional dependency of *CTT1* in the tested wild strain backgrounds.

**Figure 8.**
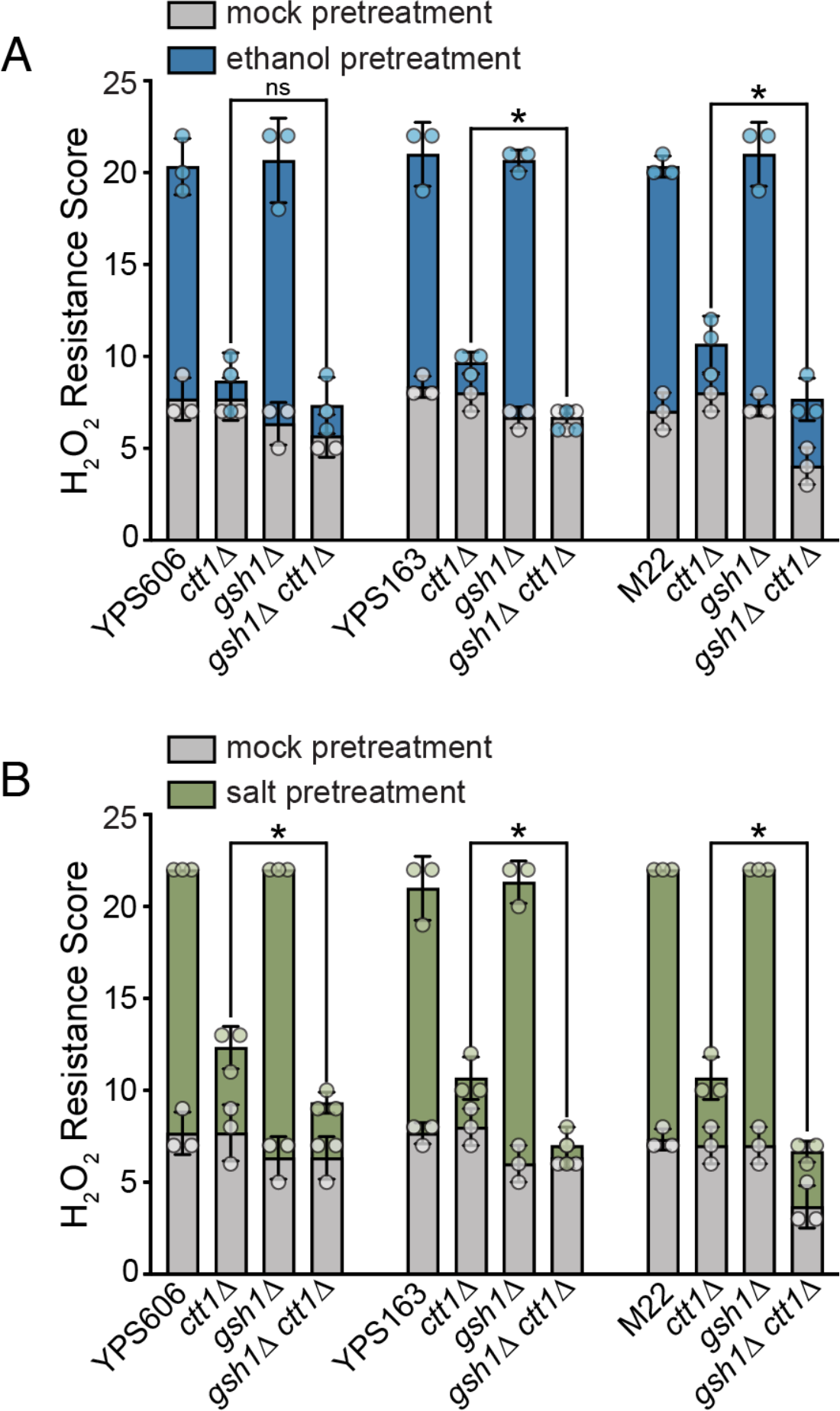
*CTT1*-dispensable acquired H2O2 resistance requires glutathione biosynthesis. Survival score plots indicating the mean and standard deviation for biological triplicates for depicted strains with either 5% ethanol (A) or 0.4 M NaCl (B) as the mild pretreatment. Error bars denote the standard deviation. The replicates for NaCl-treated YPS606, YPS606 *gsh1Δ*, M22, and M22 *gsh1*Δ strains all had the same tolerance score and thus zero standard deviation (see Tables S3 and S4 for raw numerical data). Asterisks represent significant differences in acquired resistance between denoted strains (* *P* < 0.05, ns = not significant; ordinal regression analysis on raw spot scores).

## Discussion

In this study, we leveraged natural variation in acquired stress resistance to understand the mechanistic underpinnings of this important trait. We previously showed that cytosolic catalase T (Ctt1p) was completely essential for ethanol-induced cross protection in YPS163, a wild oak strain [50]. Here, we were surprised to find substantial residual salt-induced cross protection in the YPS163 *ctt1Δ* mutant, at a level expected to be physiologically relevant [53, 54]. Moreover, about half of the strains tested showed varied levels of *CTT1* dispensability.

Interested in understanding the regulation and molecular mechanisms underlying these differences, we leveraged transcriptional profiling in strains with varied levels of *CTT1* dispensability. The results of these experiments implicated and helped identify novel regulators important for acquired stress resistance, as well as potential compensatory mechanisms that can support partial acquired H2O2 resistance in the absence of *CTT1*. This highlights the power of integrating comparative genomics with targeted mutational analysis to understand the mechanisms underlying stress defense traits.

Our analysis of regulatory mutants in the YPS606 oak strain background revealed interesting regulatory aspects for both intrinsic (basal) stress resistance and acquired stress resistance. For example, and consistent with the previous literature [59, 60], the YPS606 *yap1*Δ mutant had a severe defect in intrinsic H2O2 resistance. However, this mutant was able to fully acquire H2O2 resistance with either ethanol or salt as pretreatments, providing further support for the idea that the stress defense mechanisms underlying intrinsic and acquired stress resistance are distinct [39, 41, 42, 45, 48, 50]. The same was true of the YPS606 *skn7Δ* mutant, though its intrinsic H2O2 resistance defect was not as pronounced as that of the YPS606 *yap1*Δ mutant. In contrast, the YPS606 *msn2Δ msn4Δ* mutant had no observable defect for intrinsic H2O2 resistance, but had a strong defect for acquired H2O2 resistance when ethanol was the primary stress. Moreover, the intrinsic H2O2 resistance phenotypes of the tested transcription factor mutants correlated with differences in basal expression of genes annotated to the GO category of ‘response to oxidative stress.’ The YPS606 *msn2Δ msn4Δ* mutant, which had wild-type levels of intrinsic H2O2 resistance, also had no differentially expressed oxidative stress response genes under unstressed conditions. In contrast, the YPS606 *skn7Δ* mutant, with a moderate intrinsic H2O2 resistance defect, had four oxidative stress response genes with at least 2-fold lower basal expression than wild type: *AHP1, GPX2, PRX1*, and *TSA1*. And the YPS606 *yap1*Δ mutant, with a strong intrinsic H2O2 resistance defect, also possessed at least 2- fold lower expression of those same four genes, while also including three additional genes: *GSH1*, *HSP31*, and *SOD1*. Notable, *gsh1Δ* mutants in the M22, YPS163, YPS606, had little difference in intrinsic H2O2, suggesting that a combination of Yap1p-regulated thioredoxin- and glutathione-dependent peroxidases are responsible for intrinsic H2O2 tolerance in this particular yeast strain.

Our results also demonstrate that different yeast strains use different cellular strategies to yield superficially similar stress resistance phenotypes. For example, we showed that the paralogous general transcription factors Msn2/4p are essential in the YPS606 background for ethanol-induced cross protection against H2O2, but not salt-induced cross protection. This was somewhat surprising to us because Msn2/4p are necessary for salt-induced cross protection in the S288c background [41, 45], and highlights the importance of comparing different genetic backgrounds to obtain an integrated view of stress signaling within even a single species. While *MSN2* has been implicated in acquired ethanol (same stress) resistance [48], this is the first time it has been shown to play a role in ethanol-induced cross protection. We also found a strong defect in ethanol-induced cross protection in the YPS606 *skn7Δ* background, which has never been observed before, further highlighting the benefits of taking advantage of natural variation to identify novel biology even in well-studied model organisms [61]. Our results also provide hints about the complexity of regulation of stress responses beyond transcriptional responses. For example, YJM1129 is unable to acquire H2O2 resistance when ethanol is the primary stress, yet it had had only 1.5-fold lower relative *CTT1* mRNA abundance during ethanol stress compared to the mean of all strains. Because YJM1129 is able to acquire H2O2 resistance when salt is the primary stress, and that phenotype requires *CTT1*, it’s unlikely that allelic variation resulting in defective Ctt1p protein is responsible. One possibility is that Ctt1p protein levels are translationally regulated during stress, with low levels of translation happening during ethanol stress in the YJM1129 background. There is precedent for potential translational regulation of Ctt1p abundance during stress, as Berry and colleagues [41] found that while *CTT1* mRNA levels increase during DTT, heat shock, and NaCl stresses, only NaCl stress resulted in Ctt1p protein being detectable by western blot.

Our results implicated glutathione-dependent peroxidases as responsible for *CTT1*- independent acquisition in three tested strains that possess *CTT1*-independent acquisition. Catalases are among the most efficient and fastest-acting known enzymes [62], so it is not surprising that yeast cytosolic catalase T (Ctt1p) plays a major role in acquired H2O2 resistance. What was somewhat surprising is that some yeast strains solely rely on Ctt1p, while others supplement Ctt1p with glutathione-dependent peroxidases. One possibility is that these different strategies can benefit strains in different ecological niches. For example, since catalases use heme as a cofactor, wild strains in more-iron limited environments may supplement catalase with glutathione-dependent peroxidases, and possessing otherwise redundant catalase- independent acquired H2O2 resistance would provide a fitness advantage. On the other hand, glutathione is biosynthesized from sulfur-containing cysteine, so glutathione-dependent peroxidases may be less favored in sulfur limited conditions. Intriguingly, *CTT1* was dispensable for both ethanol and salt-induced H2O2 resistance in the vineyard strain M22, which also is known to possess resistance to copper sulfate [63], an antimicrobial agent often used in vineyards [64]. Because sulfate supplementation increases glutathione pools in yeast [65], vineyard environments in particular may favor partial glutathione-dependent strategies for H2O2 resistance. On the other hand, we cannot rule out that the possibility that *CTT1*-independent acquisition represents neutral, non-adaptive variation. Future studies with larger sample sizes are needed to identify causative loci responsible for variation in *CTT1* dispensability, and then determine whether the responsible polymorphisms are more likely to have arisen via adaptative or neutral processes.

## Conclusions

In summary, we found surprising heterogeneity across diverse yeast strains in *CTT1* dispensability for acquired H2O2 resistance following mild ethanol and/or salt pretreatment. Transcriptional profiling of ethanol and salt stress responses in strains with different levels of *CTT1* dispensability implicated potential regulators of acquired stress resistance, as well as glutathione-dependent peroxidases responsible for *CTT1*-independent acquisition. Ultimately, this study highlights how superficially similar traits can have different underlying molecular foundations and provides a framework for understanding diversity and regulation of stress defense mechanisms.

## Materials and Methods

### Strains and growth conditions

Strains and primers used in this study are listed in S1 and S2 Tables, respectively. Fully prototrophic diploids in the S288c (DBY8268) background were generated by repairing the *ura3* auxotrophic marker in both MATa (JL505) and MATa (JL506) haploids (whose construction is described in [49]) via transformation of a PCR product containing the M22 *URA3* allele, followed by mating the two prototrophic haploids to generate diploid strain JL518. Because wild yeast strains can be heterozygous, we generated isogenic homozygous diploid segregants of wild yeast strains prior to genetic manipulation. Sporulation was induced by growing cells for 48 hours in YPD (1% yeast extract, 2% peptone, 2% dextrose), harvesting by centrifugation (1,500 x *g* for 3 minutes), resuspending in 1% potassium acetate, and incubating for 2–10 days at 25°C with orbital shaking (270 rpm). Tetrads were then dissected and streaked to obtain colonies consisting of ‘selfed’ homozygous diploids (which occurs because the wild strains are homothallic and thus capable of mating type switching), and diploidy was confirmed using mating-type tester strains [66]. Deletions were introduced into the different wild strains via homologous recombination following transformation of the relevant deletion::KanMX cassette amplified from the Yeast Knockout Collection [67], with the exception of *ctt1Δ*::KanMX, which was amplified from YPS163-derived JL143 [50]. To generate double or triple mutants, a KanMX marker from one mutant was first exchanged with NatMX or HygMX via homologous recombination [68], followed by transformation with the appropriate second deletion::KanMX PCR product. All deletions were verified by diagnostic PCR that included primer pairs that annealed to the MX cassette and a sequence upstream of the deletion (verifying the ‘scar’), and primer pairs that annealed within the deleted region (verifying absence of the gene). All yeast strains were grown at 30°C with orbital shaking (270 rpm). When used for selection, antibiotic concentrations were 100 µg/ml for nourseothricin (Nat), 200 µg/ml for G418 (Kan), and 300 µg/ml for hygromycin B (Hyg).

### Cross protection assays

Cross protection assays were performed as described [50]. Briefly, cells were streaked to generate isolated colonies, and 5 unique colonies per replicate experiment were inoculated and grown to saturation in YPD (OD600 ∼20 on a Unico spectrophotometer). Cultures were then sub-cultured for at least 8 generations (> 12 hours) to mid-exponential phase (OD600 0.3 – 0.6). The cultures were then split, with one sample receiving a mock treatment (diluted 1:1 into fresh media) and the other sample receiving a mild ‘primary’ dose of stress (diluted 1:1 into fresh media containing either 10% v/v EtOH (5% final concentration) or 0.8 M NaCl (0.4 M final concentration)), with ‘mild’ stress being defined as a dose that can increase survival to severe stress while not affecting viability itself (>95% survival). The mild doses of 5% EtOH and 0.4 M NaCl were initially empirically determined for a subset of strains used in this study [50], and were verified during initial experiments to not cause loss of viability across the entire panel strains (see Figure S1, comparing the leftmost control spots for mock and pretreated samples).

Following a 1-hour incubation at 30°C with orbital shaking (270 rpm), mock and pretreated cells were collected by mild centrifugation at 1,500 x *g* for 3 min, resuspended in fresh media to an OD600 of 0.6, and then diluted 3-fold (150 µl total volume) into a microtiter plate containing a panel of 11 severe ‘secondary’ H2O2 doses ranging from 0.75 – 5 mM (0.75 mM, 1 mM, 1.25 mM, 1.5 mM, 1.75 mM, 2 mM, 2.5 mM, 3 mM, 3.5 mM, 4 mM, 5 mM). Microtiter plates were sealed with breathable Rayon films (VWR), and cells were incubated with secondary stress for 2 hours at 30°C with 800 rpm shaking in a VWR Symphony incubating microplate shaker. Cells were diluted 50-fold in YPD a microtiter plate, and then 4 µl of each well was spotted onto YPD agar plates and grown for 48 hours at 30°C. Viability at each dose was scored using a 4-point semi-quantitative scale comparing viability to a no secondary stress (YPD only) control: 100% viability = 3 pts, 50–90% viability = 2 pts, 10–50% viability = 1 pt, and <10% (3 or fewer colonies) = 0 pts. An overall acquired resistance H2O2 score was calculated as the sum of the raw spot scores for each of the 11 doses of secondary stress. Raw tabular data used to generate figures can be found in Tables S3 and S4. A detailed acquired stress resistance protocol can be found on protocols.io under doi dx.doi.org/10.17504/protocols.io.g7sbzne.

Experiments where statistical analysis was performed were done in at least triplicate. For experiments shown in Figures 1 and Figure 8, replicates for each batch of strains were performed on separate days. For the transcription factor mutant experiments shown in Figures 5 and S2, there were too many samples to run as a single batch. Instead, each transcription factor mutant being tested was run alongside a wild-type and *ctt1Δ* mutant control, resulting in triplicate values for each set of transcription factor mutants and 24 replicates for the wild-type and *ctt1Δ* control strains. While bar graphs depict the mean and standard deviations for overall resistance scores, relevant statistical comparisons were performed on the raw spot score data. Because the spot score data is ordinal with spacing that cannot be assumed to be equidistant, we used the proportional-odds cumulative logit model assess statistical significance of differences in stress resistance for comparisons of interest depicted in Figures 1, 5, and 8, with Benjamini-Hochberg corrected *P*-values being reported. The proportional-odds cumulative logit model is a type of ordinal regression that is used to model the relationship between an ordinal response variable (i.e., resistance as measured by the raw spot scores) and one or more predictor variables (i.e., genotype and/or pretreatment) [69], and has been used in ecological and biomedical contexts to analyze ordinal data [70, 71]. A detailed summary of the approach and outputs is described in File S1, and an R Markdown file with the code used for analyses is provided in File S2.

For classifying strains according to *CTT1* dispensability, percent residual H2O2 acquisition was calculated using the summed tolerance scores as: *ctt1*Δ mild stress pretreatment score – *ctt1*Δ mock pretreatment score WT mild stress pretreatment score – WT mock pretreatment score Strains with at least 10% residual acquisition were categorized as *CTT1* being dispensable for a given mild stress pretreatment. Hierarchical clustering of tolerance scores was performed using Cluster 3.0 (http://bonsai.hgc.jp/~mdehoon/software/cluster/software.htm), using Euclidean distance as the similarity metric and centroid linkage with a weighted cutoff of 0.4 and an exponent value of 1 [72], and visualized using Java TreeView (https://sourceforge.net/projects/jtreeview/files/) [73].

### RNA sequencing and analysis

Two separate RNA-seq experiments were performed: i) comparison of phenotypically diverse strains (M1, M22, S288c, Y10, YJM1129, YJM308, YPS163, YPS606) responding to ethanol or NaCl stress relative to an unstressed control (performed in biological triplicate, except for the Y10 ethanol-stressed samples, where one sample was found post-analysis to be a mix-up) and ii) comparison of YPS606 vs. YPS606 transcription factor mutants (*msn2Δ msn4Δ*, *skn7Δ*, *yap1Δ*) responding to ethanol or NaCl stress relative to an unstressed control (performed in biological quadruplicate). For both sets of experiments, cells were grown at least 8 generations in 80 ml YPD to mid-exponential phase (OD600 0.3 – 0.6). Fifteen-ml of an unstressed control sample was taken, and then the culture was split, with 20 ml of culture being added to 20-ml media containing either 10% v/v ethanol (5% final concentration) or 0.8 M NaCl (0.4 M NaCl final concentration). Following incubation in stress media for either 30 minutes (ethanol) or 45 minutes (NaCl), 15 ml of cells were collected by centrifugation at 1,500 x *g* for 1 minute, flash frozen in liquid nitrogen, and stored at -80°C until processed.

RNA was extracted using the ‘hot phenol’ method [74], and a detailed protocol is available at protocols.io under DOI dx.doi.org/10.17504/protocols.io.inwcdfe. Phenol-extracted total RNA was treated with DNase I (Ambion), and then purified with a Quick-RNA MiniPrep Plus Kit (Zymo Research) including the optional on-column DNase digestion. RNA integrity for each sample was assessed using an Agilent 2200 TapeStation, and all samples had a RIN value above 9.0.

RNA-seq libraries were prepared using the KAPA mRNA HyperPrep Kit (Roche) and KAPA Single-Indexed Adapter Set A + B, according to manufacturer specifications with minor modifications. Briefly, 500 ng of total RNA was used as starting material for each sample, polyA-enrichment was performed, mRNA was fragmented to an average size of 200–300 nucleotides, and the cDNA libraries were amplified for 9 PCR cycles. A detailed a protocol is available on protocols.io under DOI dx.doi.org/10.17504/protocols.io.uueewte. cDNA libraries were pooled (24-plex) and sequenced on a HiSeq4000 at the University of Chicago Genomics Facility, generating 50-bp single-end reads. To minimize batch effects, a random block design was used for library construction for each biological replicate, and each biological replicate was sequenced together on a single lane.

Reads were trimmed of low-quality reads and adapter sequences using Trimmomatic (version 0.38) [75], with the following commands: ILLUMINACLIP:Kapa_indices.fa:2:30:10 LEADING:3 TRAILING:3 MAXINFO:40:0.4 MINLEN:40. To generate new reference genomes for each wild strain, RNA-seq reads for each individual strain were first mapped to the S288c genome (version Scer3) using Bowtie 2 (version 2.3.4.1) [76], variants were called using bcftools (version 1.9) [77], and then variant calls were used to generate new reference genomes using GATK3 (Version 3.8-1-0) [78]. Trimmed reads were then mapped to their new corresponding reference genomes using STAR (version 2.6.1) [79]. Expected counts for each transcript were generated using RSEM (version 1.3.1) [80], and are reported in Table S5.

Differential expression analysis was performed using edgeR (version 3.26.4), using TMM normalization on all genes with at least one count per million (CPM) in at least one strain-condition, and the quasi-likelihood (QL) model [81], using sample type (e.g., S288c unstressed, S288c ethanol stressed, S288c NaCl stressed…) and biological replicate as factors.

To identify strain-specific differences in stress responses, the response of each strain was compared to the mean response of all strains. To identify sub-classes of strain-specific responses, we generated a ‘consensus’ response to each stress by using only treatment (unstressed, ethanol stressed, or NaCl stressed) and biological replicate as factors. Genes with significant differential expression (FDR < 0.01) of at least ± 1.5-fold were considered to be induced or repressed by each stress. Then for each strain comparison, genes with significant differential expression (FDR < 0.01) of at least ± 1.5-fold compared to the mean of all strains were classified based on the direction of the difference (e.g., genes with >1.5-fold higher expression compared to the mean of all strains for ‘consensus’ induced genes were classified as having “amplified induction,” while a sign shift from ‘consensus’ induction to >1.5-fold repression resulted in genes being classified as “novel repression”). All raw RNA-seq data are available through the National Institutes of Health Gene Expression Omnibus (GEO) database under accession no. GSE248219, and the edgeR outputs can be found in Table S6.

Hierarchical clustering of gene expression log2 fold changes was performed with Cluster 3.0, using Euclidean distance as the similarity metric, centroid linkage with a weighted cutoff of 0.4 and an exponent value of 1, and visualized in Java TreeView. Functional enrichments of gene ontology (GO) categories were performed using Princeton’s GO-TermFinder (https://go.princeton.edu/cgibin/GOTermFinder) [82], with Bonferroni-corrected *P*-values < 0.01 taken as significant. Significantly enriched regulatory associations (FDR-corrected *P* < 0.01) were identified using the YEAst Search for Transcriptional Regulators And Consensus Tracking (YEASTRACT) database [83], using documented DNA binding or expression evidence (with the transcription factor acting as an activator or unknown), filtered on conditions defined as stress. GO and regulatory enrichments can be found in Table S7.

## Acknowledgements

We thank Dr. Justin Fay and Dr. Audrey Gasch for wild yeast strains, Dr. Elizabeth Ruck for running samples on the TapeStation, and Dr. Andrew Alverson for the use of equipment. This work was supported in part by National Science Foundation Grants IOS-1656602 (JAL) and MCB-1941824 (JAL), the Arkansas Biosciences Institute (Arkansas Settlement Proceeds Act of 2000) (JAL), and a Research Assistantship provided through the University of Arkansas Cell and Molecular Biology Graduate Program (ANS). The funders had no role in study design, data collection and interpretation, or the decision to submit the work for publication.

## Availability of data and materials

All raw data generated and analyzed for this study are included in either the supplementary files or in public data repositories. Raw RNA-seq read data are available through the National Institutes of Health Gene Expression Omnibus (GEO) database under accession no. GSE248219. Raw tabular data used to generate figures can be found in Tables S3 and S4.

## Supporting Figures

**Figure S1.**
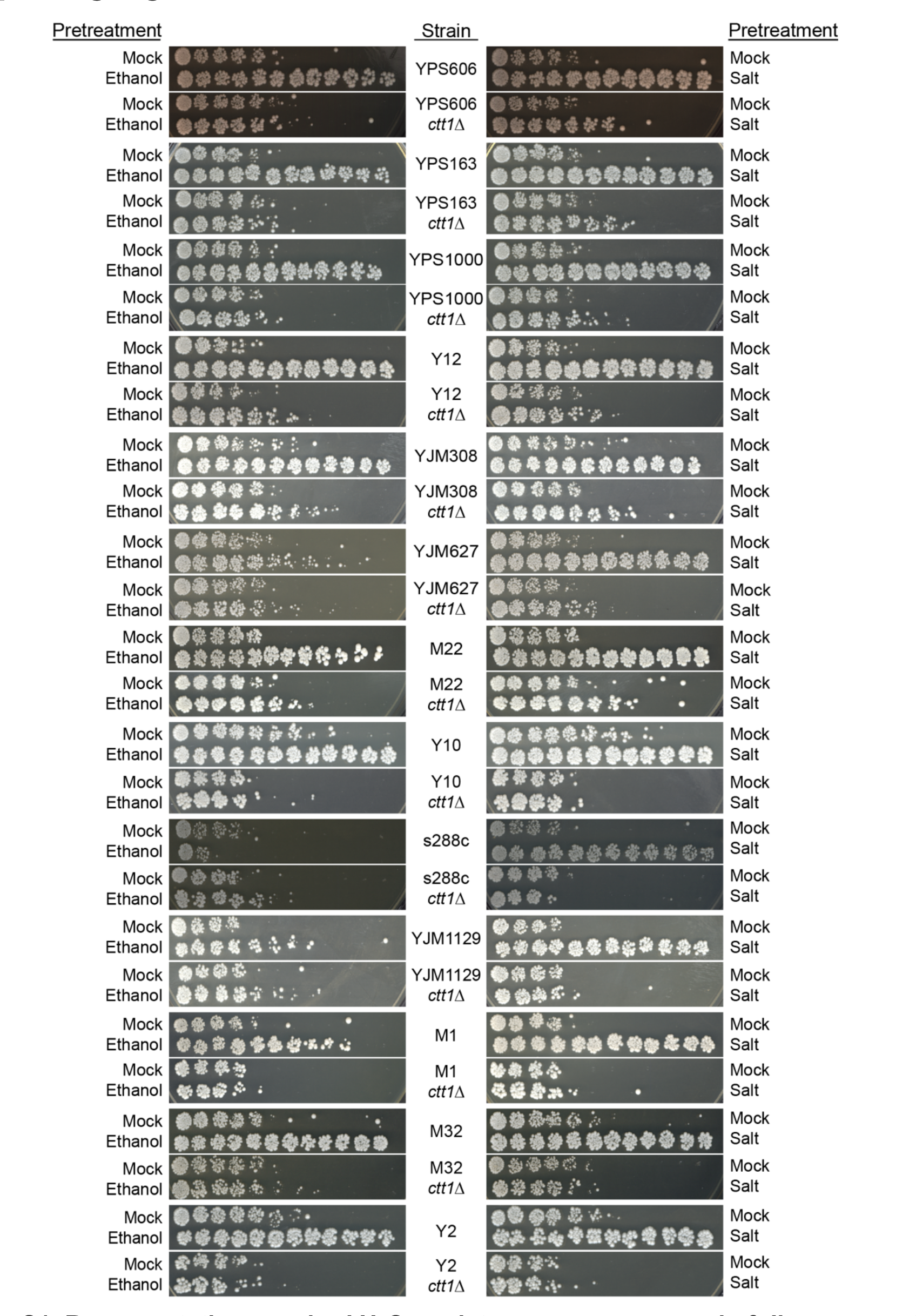
Representative acquired H2O2 resistance assays a panel of diverse yeast strains. Representative acquired H2O2 resistance assays are shown for all strains depicted in Figure 2.

**Figure S2.**
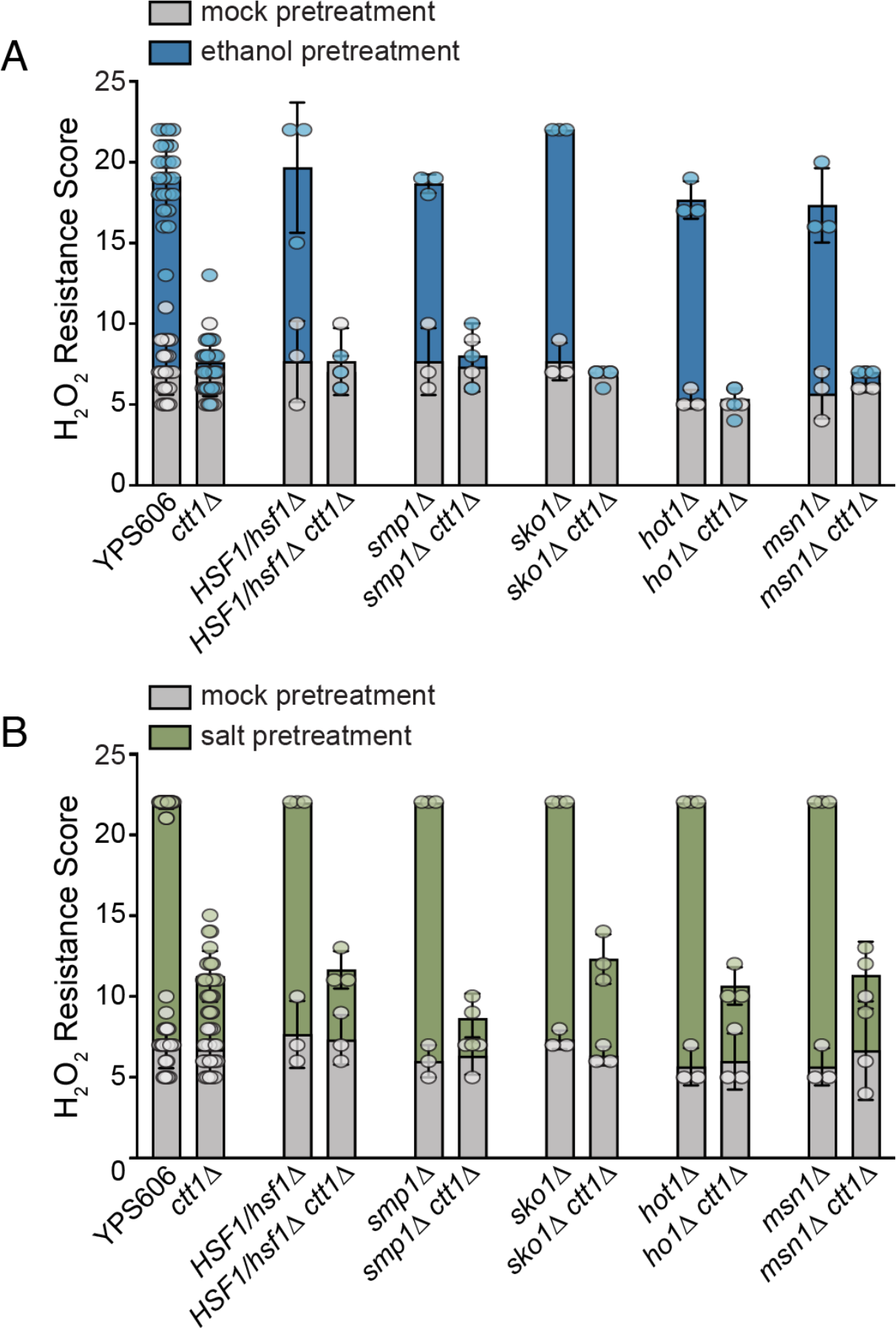
Acquired H2O2 resistance assays for a panel YPS606 transcription factor deletion mutants. Acquired H2O2 resistance assays were performed in homozygous YPS606 *sko1Δ*, *hot1Δ*, and *msn1Δ* mutants plus a heterozygous *HSF1/hsf1Δ* mutant (because *HSF1* is essential). Mutants were assayed in biological triplicate, while wild-type YPS606 and the *ctt1Δ* single mutant were included as controls for each set of experiments and thus had 24 replicates each. Error bars depict the standard deviation (note that the replicates some strain comparisons had the exact same tolerance score and thus zero standard deviation).

**Figure S3.**
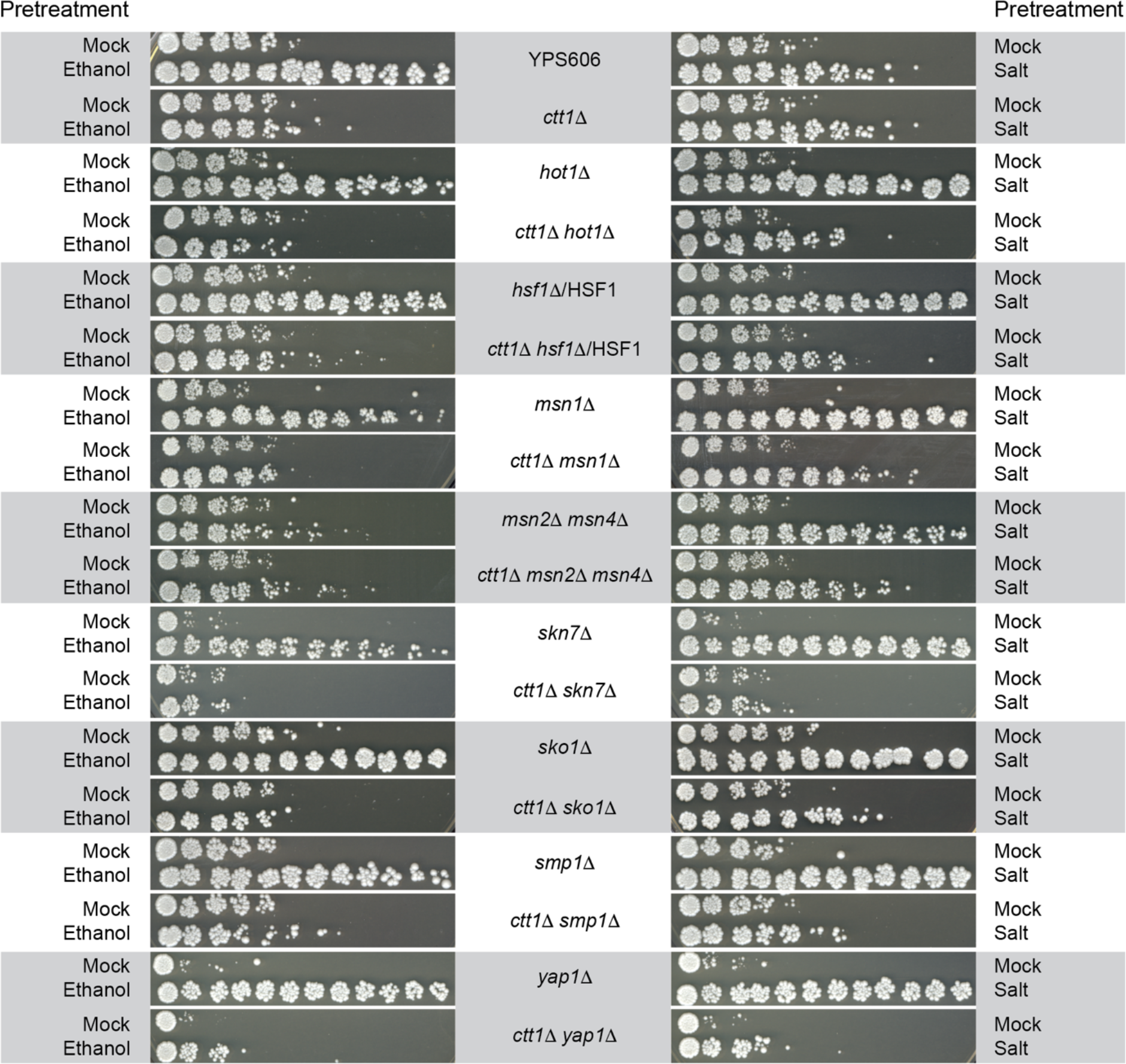
Representative acquired H2O2 resistance assays for all transcription factor mutants. Representative acquired H2O2 resistance assays are shown for all strains depicted in Figures 6 and S2.

**Figure S4.**
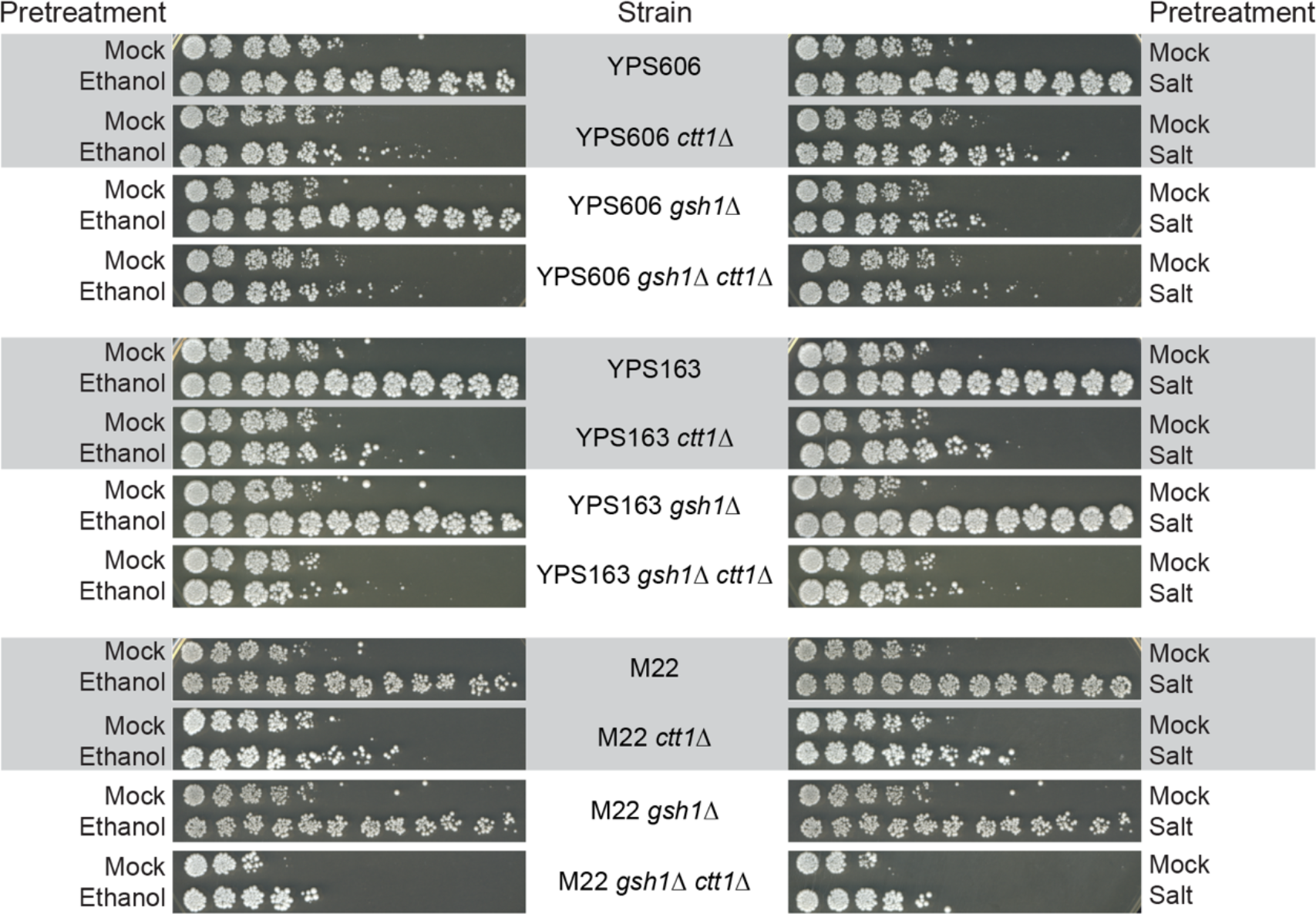
Representative acquired H2O2 resistance assays for YPS606 *gsh1Δ* mutants. Representative acquired H2O2 resistance assays are shown for all strains depicted in Figure 8.

## Supporting Tables and Files

Table S1: Strains used in this study.

Table S2: Primers used in this study.

Table S3: Raw data used to generate figures.

Table S4: Ordinal data used for statistical analyses.

Table S5: RSEM Output.

Table S6: edgeR Output.

Table S7: GO and TF binding-site enrichments for clusters in Figure 3.

Table S8: Categories of differential ethanol-responsive expression in Figure 4.

Table S9: Categories of differential salt-responsive expression in Figure 4.

File S1: Supporting information on ordinal analysis of stress viability assays.

File S2: R Markdown file for ordinal analysis of stress viability assays.

## Notes

### Competing Interest Statement

The authors have declared no competing interest.

### Summary of Updates

The manuscript was revised to include a better conceptual framework for "many-to-one mapping" applied to stress defense. Ordinal statistical analyses were applied to semi-quantitative data, and the Methods and Results including Figures 1, 5, and 8 were updated. New authors who performed the statistical analyses were added. Supplemental files were updated to include the raw ordinal data, a description of the statistical models, and code for performing the analyses.

