## Supplementary material for "Natural variation in yeast reveals multiple paths for acquiring higher stress resistance": File S1

### Ordinal Analysis of Semi-Quantitative Stress Viability Data

The raw spot scores for viability in stress resistance assays are on a semi-quantitative ordinal scale, and are then summed to generate an overall resistance score. There is debate about whether summed ordinal data can be appropriately analyzed via ‘classic’ parametric statistical tests such as t-tests [1]. Non-parametric tests such as the Wilcoxon rank-sum test would be appropriate, but are underpowered for biological triplicates. However, an ordinal regression model is appropriate for analyzing raw ordinal data that has a natural ordering (0 < 1 < 2 < 3) and spacing that cannot be assumed to be equidistant [2, 3]. The proportional-odds cumulative logit model is a type of ordinal regression model that is used to model the relationship between an ordinal response variable (e.g., level of stress resistance) and one or more predictor variables (e.g., pretreatment and/or strain genotype) [4], which can be used to assess statistical significance for comparisons of interest.

The models used below assume that the log-odds of the response variable being less than or equal to a given category is a function of the predictor variable genotype and/or pretreatment. This model is a generalization of the binary logistic regression model to the case of more than two levels of the response. This generalized linear model (GLM) approach allows us to model the relationship between the genotype and/or pretreatment on the raw ordinal semi-quantitative scores. Additionally, this model avoids the issues that arise treating ordinal values as specific values. As with linear regression, the proportional-odds cumulative logit model estimates the effect of changes in the predictor variables on the response variable, through the link function. Let $Y$ be the score with categories $j = 0,1,2$, resulting in $J = 3$, where $J$ is the number of categories with non-zero probabilities. The model structure for comparing genotypes is given by:

$$log\frac{P(Y \leq j)}{P(Y > j)} = \alpha_j + \beta_{gene\Delta}X_{gene\Delta} + \beta_{[H_2O_2]}X_{[H_2O_2]}$$
$$log\frac{P(Y \leq j)}{P(Y > j)} = \alpha_j + \beta_{prestress}X_{prestress} + \beta_{[H_2O_2]}X_{[H_2O_2]}$$

where the model is a system of $J - 1$ equations, one for each cumulative logit (e.g., $P(y_i \leq 0))$, $P(y_i \leq 1)$ for $j \in \{0,1\}$. To interpret, note that $\alpha_0$ is the estimated log-odds of a score of $0$ when all predictors are their baseline, that is: $X_{gene\Delta} = X_{[H_2O_2]} = 0$. Similarly, $\alpha_1$ is the estimated log odds of a score of $1$ or $0$ when all predictors are at their baseline. The parameter estimates of greater interest is $\hat{\beta}_{gene\Delta}$, the estimated effect on the cumulative log odds ratio of a given gene deletion compared to its baseline genotype. When comparing WT vs *ctt1*Δ as in Figure 1, the baseline category is WT. In comparisons of *ctt1*Δ vs *ctt1*Δ*gsh1*Δ , then *ctt1*Δ is coded as the baseline genotype. $\hat{\beta}_{prestress}$ is the estimated main effect of mild stress pretreatment relative to mock pretreatment. This model is used when comparing the effect of mild stress pretreatment on isogenic samples. $\hat{\beta}_{[H_2O_2]}$ is the estimated effect of an increase in 1 mM concentration of peroxide stress on cumulative log-odds ratio of the survival score. The link function for this model is the cumulative logit function, which is the natural logarithm of the odds of the response variable being less than or equal to a given category.

A powerful motivation for this model is its connection to the idea of a continuous latent response. Suppose an undetected but existing continuous variable $Z$ (e.g. “stress sensitivity”) exists, and $Z$ is described by the measured ordinal score $Y$, then the proportional odds cumulative logit model is similar to that of a linear regression on $Z$ with a logistic error function.

The relevant model above was fit for each comparison shown below. The proportional odds assumption was satisfied in many models, assessed by deviance difference test $(p > 0.05)$. For those for which it was not, a non-proportional odds model was fit and results matched between the proportional odds model were found to be significant in at least one level for the non-proportional odds model. The proportional odds model was used to estimate effects for these data. As suggested by Agresti [5], if it is plausible to fit an ordinary linear regression model fitting the latent variable measured by the ordinal score response, then the assumption of proportional odds is sensible. Furthermore, the effects of interest are on the aggregate effect across all score levels of the response, which is provided by the proportional odds model. In addition, we also fit adjacent category non-proportional odds models to data that may violate the assumption. The outputs of these models are available in this supplement. While these models by design provide estimates for each transition between ordinal levels, results were highly comparable. For clarity and consistency, the results from proportional odds model fits from VGAM::vlgm are used in the figures of the manuscript.

#### Estimated Coefficients of Interest

Fig 1: Noted Comparisons

| Strain | Treatment | Comparison | Estimate | std.error | statistic | P-Value | FDR[1] |
|---|---|---|---|---|---|---|---|
| S288c | EtOH | WTvsCtt1 | −6.68 | 1.99 | −3.35 | 8.15e-04 | 0.006 |
| YPS163 | NaClvsMock | Ctt1 | −4.30 | 1.28 | −3.36 | 7.74e-04 | 0.006 |

*[1] Benjamini-Hochberg calculated false discovery rate (FDR) [6]*

Fig 5: Noted Comparisons

| Strain | Treatment | Comparison | Estimate | std.error | statistic | P-Value | FDR |
|---|---|---|---|---|---|---|---|
| YPS606 | EtOH | WTvsMsn24 | 3.27 | 0.434 | 7.54 | 4.59e-14 | 6.42e-13 |
| YPS606 | EtOH | WTvsSkn7 | 2.11 | 0.405 | 5.22 | 1.76e-07 | 4.94e-07 |
| YPS606 | EtOH | Ctt1vsCtt1Yap1 | 5.22 | 0.931 | 5.61 | 2.04e-08 | 7.14e-08 |
| YPS606 | EtOH | Ctt1vsCtt1Skn7 | 3.47 | 0.803 | 4.32 | 1.56e-05 | 3.64e-05 |
| YPS606 | NaCl | Ctt1vsCtt1Yap1 | 6.46 | 0.928 | 6.96 | 3.39e-12 | 2.37e-11 |
| YPS606 | NaCl | Ctt1vsCtt1Skn7 | 4.88 | 0.803 | 6.08 | 1.19e-09 | 5.57e-09 |

Fig 8: Noted Comparisons

| Strain | Treatment | Comparison | Estimate | std.error | statistic | P-Value | FDR |
|---|---|---|---|---|---|---|---|
| YPS606 | EtOH | Ctt1vsCtt1Gsh1 | 1.74 | 1.04 | 1.67 | 0.094 | 0.220 |
| YPS163 | EtOH | Ctt1vsCtt1Gsh1 | 3.40 | 1.14 | 2.97 | 0.003 | 0.015 |
| M22 | EtOH | Ctt1vsCtt1Gsh1 | 2.10 | 0.815 | 2.57 | 0.010 | 0.028 |
| YPS606 | NaCl | Ctt1vsCtt1Gsh1 | 5.49 | 2.08 | 2.64 | 0.008 | 0.028 |
| YPS163 | NaCl | Ctt1vsCtt1Gsh1 | 5.15 | 1.75 | 2.95 | 0.003 | 0.015 |
| M22 | NaCl | Ctt1vsCtt1Gsh1 | 3.17 | 1.00 | 3.15 | 0.002 | 0.015 |

#### Testing Proportional Odds Assumption

The null hypothesis for this deviance difference test suggest that the proportional odds assumption is consistent with the data. The assumption is supported in some comparisons and inconsistent with the data in others. However, the underlying logic of a continuous latent variable being approximated by the semi-quantitative scores is reasonable and more easily interpreted. As such, values reported here are for models with proportional odds.

Tests of Proportional Odds Assumption

| Figure | Strain | Pretreatment | Genotype | P-Value |
|---|---|---|---|---|
| 1 | S288c | EtOH | WT vs ctt1 | 0.895 |
| 1 | YPS163 | NaCl vs Mock | ctt1 | 0.039 |
| 5 | YPS606 | EtOH | msn2msn4 vs WT | 0.099 |
| 5 | YPS606 | EtOH | skn7 vs WT | 0.164 |
| 5 | YPS606 | EtOH | ctt1 skn7 vs ctt1 | 0.338 |
| 5 | YPS606 | EtOH | ctt1 yap1 vs ctt1 | 0.141 |
| 5 | YPS606 | NaCl | ctt1 skn7 vs ctt1 | 0.007 |
| 5 | YPS606 | NaCl | ctt1 yap1 vs ctt1 | 0.001 |
| 8 | YPS163 | NaCl | ctt1 gsh1 vs ctt1 | $4.9 \times 10^{-5}$ |
| 8 | YPS606 | NaCl | ctt1 gsh1 vs ctt1 | 0.650 |
| 8 | M22 | NaCl | ctt1 gsh1 vs ctt1 | $9.5 \times 10^{-8}$ |
| 8 | YPS163 | EtOH | ctt1 gsh1 vs ctt1 | 0.028 |
| 8 | YPS606 | EtOH | ctt1 gsh1 vs ctt1 | 0.970 |
| 8 | M22 | EtOH | ctt1 gsh1 vs ctt1 | $2.6 \times 10^{-7}$ |

#### Ensuring significance without proportional odds assumption

For models for which there was evidence of non-proportional odds, a cumulative logit model without a proportional odds assumption was fit and estimates for the contrast of interest were extracted. In general, the system of equations for the non-proportional odds is:

$$log\frac{P(Y \leq j)}{P(Y > j)} = \alpha_j + \beta_{j,gene\Delta}X_{gene\Delta} + \beta_{j,[H_2O_2]}X_{[H_2O_2]}$$

$$log\frac{P(Y \leq j)}{P(Y > j)} = \alpha_j + \beta_{j,prestress}X_{prestress} + \beta_{j,[H_2O_2]}X_{[H_2O_2]}$$

Note that the effects of $X$ values are allowed to differ for different logits in this model. To illustrate, the system of equations for the cumulative logit model comparing Figure 1's S288c during EtOH pretreatment comparing WT vs *ctt1*Δ, with the proportional odds model:

$$\begin{aligned} log\frac{P(Y \leq 0)}{P(Y > 0)} &= \alpha_0 + \beta_{gene\Delta}X_{gene\Delta} + \beta_{[H_2O_2]}X_{[H_2O_2]} \\ &= -7.49 - 6.68X_{ctt1\Delta} + 8.36X_{[H_2O_2]} \\ log\frac{P(Y \leq 1)}{P(Y = 2)} &= \alpha_1 + \beta_{gene\Delta}X_{gene\Delta} + \beta_{[H_2O_2]}X_{[H_2O_2]} \\ &= -5.04 - 6.68X_{ctt1\Delta} + 8.36X_{[H_2O_2]} \end{aligned}$$

and the non-proportional odds model:

$$log\frac{P(Y \leq 0)}{P(Y > 0)} = \alpha_0 + \beta_{\mathbf{0},gene\Delta}X_{gene\Delta} + \beta_{\mathbf{0},[H_2O_2]}X_{[H_2O_2]}$$
$$= -6.70 - 6.45X_{ctt1\Delta} + 7.67X_{[H_2O_2]}$$
$$log\frac{P(Y \leq 1)}{P(Y = 2)} = \alpha_1 + \beta_{\mathbf{1},gene\Delta}X_{gene\Delta} + \beta_{\mathbf{1},[H_2O_2]}X_{[H_2O_2]}$$
$$= -5.61 - 6.10X_{ctt1\Delta} + 8.47X_{[H_2O_2]}$$

Note that the estimated effects for genotype and hydrogen peroxide can vary between each logit in the system of equations, with positive coefficients, as seen here for hydrogen peroxide, corresponding to an increased probability of the sample belonging to a lower score as that covariate increases. The estimates for $\beta_j$ for all comparisons for which the proportional odds assumption is statistically violated are plotted below. Red corresponds to a 95% confidence interval (CI) that does not contain 0 and blue to non-significance. Note that the only comparison for which no estimate is nonzero is the YPS606 comparison from Figure 8A, which was also non-significant in the proportional-odds model estimate shown above. This line of evidence supports the robustness of our statistical interpretations.

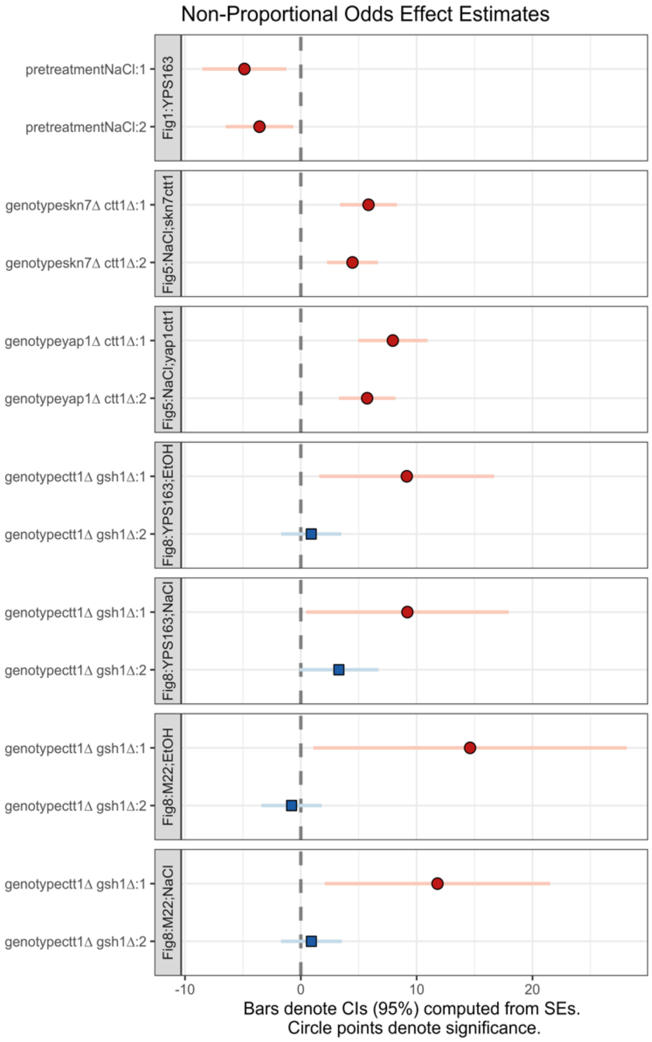

Analyses were conducted using the R Statistical language (version 4.3.1; R Core Team, 2023) on macOS Ventura 13.6.7, using the packages gt (version 0.10.1; Iannone R et al., 2024), texreg (version 1.39.3; Leifeld P, 2013), report (version 0.5.8; Makowski D et al., 2023), here (version 1.0.1; Müller K, 2020), readxl (version 1.4.3; Wickham H, Bryan J, 2023), dplyr (version 1.1.2; Wickham H et al., 2023), purrr (version 1.0.1; Wickham H, Henry L, 2023), tidyr (version 1.3.0; Wickham H et al., 2023) and VGAM (version 1.1.10; Yee TW, 2015).


- Iannone R, Cheng J, Schloerke B, Hughes E, Lauer A, Seo J (2024). *gt: Easily Create Presentation-Ready Display Tables*. R package version 0.10.1, https://CRAN.R-project.org/package=gt.
- Leifeld P (2013). “texreg: Conversion of Statistical Model Output in R to LaTeX and HTML Tables.” *Journal of Statistical Software*, *55*(8), 1-24. https://doi.org/10.18637/jss.v055.i08.
- Makowski D, Lüdecke D, Patil I, Thériault R, Ben-Shachar M, Wiernik B (2023). “Automated Results Reporting as a Practical Tool to Improve Reproducibility and Methodological Best Practices Adoption.” *CRAN*. https://easystats.github.io/report/.
- Müller K (2020). *here: A Simpler Way to Find Your Files*. R package version 1.0.1, https://CRAN.R-project.org/package=here.
- R Core Team (2023). *R: A Language and Environment for Statistical Computing*. R Foundation for Statistical Computing, Vienna, Austria. https://www.R-project.org/.
- Wickham H, Bryan J (2023). *readxl: Read Excel Files*. R package version 1.4.3, https://CRAN.R-project.org/package=readxl.
- Wickham H, François R, Henry L, Müller K, Vaughan D (2023). *dplyr: A Grammar of Data Manipulation*. R package version 1.1.2, https://CRAN.R-project.org/package=dplyr.
- Wickham H, Henry L (2023). *purrr: Functional Programming Tools*. R package version 1.0.1, https://CRAN.R-project.org/package=purrr.
- Wickham H, Vaughan D, Girlich M (2023). *tidyr: Tidy Messy Data*. R package version 1.3.0, https://CRAN.R-project.org/package=tidyr.
- Yee TW (2015). *Vector Generalized Linear and Additive Models: With an Implementation in R*. Springer, New York, USA. Yee TW, Wild CJ (1996). “Vector Generalized Additive Models.” *Journal of Royal Statistical Society, Series B*, *58*(3), 481-493. Yee TW (2010). “The VGAM Package for Categorical Data Analysis.” *Journal of Statistical Software*, *32*(10), 1-34. doi:10.18637/jss.v032.i10 https://doi.org/10.18637/jss.v032.i10. Yee TW, Hadi AF (2014). “Row-column interaction models, with an R implementation.” *Computational Statistics*, *29*(6), 1427-1445. Yee TW (2024). *VGAM: Vector Generalized Linear and Additive Models*. R package version 1.1-10, https://CRAN.R-project.org/package=VGAM. Yee TW (2013). “Two-parameter reduced-rank vector generalized linear models.” *Computational Statistics and Data Analysis*, *71*, 889-902. Yee TW, Stoklosa J, Huggins RM (2015). “The VGAM Package for Capture-Recapture Data Using the Conditional Likelihood.” *Journal of Statistical Software*, *65*(5), 1-33. doi:10.18637/jss.v065.i05 https://doi.org/10.18637/jss.v065.i05. Yee TW (2020). “The VGAM package for negative binomial regression.” *Australian and New Zealand Journal of Statistics*, *62*(1), 116-131.
