## Supplementary material for "Natural variation in yeast reveals multiple paths for acquiring higher stress resistance": File S2

---
title: "Proportional-Odds Cumulative Logit Model Analysis "
author: "Carson Stacy, Qingyang Zhang, and Jeffrey Lewis"
date: "`r Sys.Date()`"
output:
  pdf_document: default
  html_document:
    df_print: paged
  word_document: default
bibliography: references.bib
---

```{r setup, include=FALSE}
knitr::opts_chunk$set(echo = TRUE, warning = FALSE, echo=FALSE)
```

# Ordinal Analysis of Semi-Quantitative Stress Viability Data

The raw spot scores for viability in stress resistance assays are on a semi-quantitative ordinal scale, and are then summed to generate an overall resistance score. There is debate about whether summed ordinal data can be appropriately analyzed via 'classic' parametric statistical tests such as t-tests @norman2010. Non-parametric tests such as the Wilcoxon rank-sum test would be appropriate, but are underpowered for biological triplicates. However, an ordinal regression model is appropriate for analyzing raw ordinal data that has a natural ordering (0 \< 1 \< 2 \< 3) and spacing that cannot be assumed to be equidistant @lee1992 @guisan2000. The proportional-odds cumulative logit model is a type of ordinal regression model that is used to model the relationship between an ordinal response variable (e.g., level of stress resistance) and one or more predictor variables (e.g., pretreatment and/or strain genotype) @mccullagh1980, which can be used to assess statistical significance for comparisons of interest.

The models used below assume that the log-odds of the response variable being less than or equal to a given category is a function of the predictor variable genotype and/or pretreatment. This model is a generalization of the binary logistic regression model to the case of more than two levels of the response. This generalized linear model (GLM) approach allows us to model the relationship between the genotype and/or pretreatment on the raw ordinal semi-quantitative scores. Additionally, this model avoids the issues that arise treating ordinal values as specific values. As with linear regression, the proportional-odds cumulative logit model estimates the effect of changes in the predictor variables on the response variable, through the link function. Let $Y$ be the score with categories $j = 0,1,2$, resulting in $J=3$, where $J$ is the number of categories with non-zero probabilities. The model structure for comparing genotypes is given by:

$$
\begin{split}
log{\frac{P(Y\le j)}{P(Y>j)}} &= \alpha_j + \beta_{gene\Delta}X_{gene\Delta} + \beta_{[H_2O_2]}X_{[H_2O_2]} \\
log{\frac{P(Y\le j)}{P(Y>j)}} &= \alpha_j + \beta_{prestress}X_{prestress} + \beta_{[H_2O_2]}X_{[H_2O_2]}
\end{split}
$$

where the model is a system of $J-1$ equations, one for each cumulative logit (e.g., $P(y_i \le 0)),\ P(y_i \le 1)$ for $j \in \{0,1\}$. To interpret, note that $\alpha_0$ is the estimated log-odds of a score of $0$ when all predictors are their baseline, that is: $X_{gene\Delta}=X_{[H_2O_2]}=0$. Similarly, $\alpha_1$ is the estimated log odds of a score of $1$ or $0$ when all predictors are at their baseline. The parameter estimates of greater interest is $\hat\beta_{gene\Delta}$, the estimated effect on the cumulative log odds ratio of a given gene deletion compared to its baseline genotype. When comparing $ctt1\Delta$ vs $WT$ as in Figure 1, the baseline category is $WT$. In comparisons of $ctt1\Delta$ vs $gsh1\Delta\ ctt1\Delta$, then $ctt1\Delta$ is coded as the baseline genotype. $\hat\beta_{prestress}$ is the estimated main effect of mild stress pretreatment relative to mock pretreatment. This model is used when comparing the effect of mild stress pretreatment on isogenic samples. $\hat\beta_{[H_2O_2]}$ is the estimated effect of an increase in $1\ mM$ concentration of peroxide stress on cumulative log-odds ratio of the survival score. The link function for this model is the cumulative logit function, which is the natural logarithm of the odds of the response variable being less than or equal to a given category.

A powerful motivation for this model is its connection to the idea of a continuous latent response. Suppose an undetected but existing continuous variable $Z$ (e.g. "stress sensitivity") exists, and $Z$ is described by the measured ordinal score $Y$, then the proportional odds cumulative logit model is similar to that of a linear regression on $Z$ with a logistic error function. The relevant model above was fit for each comparison shown. The proportional odds assumption was satisfied in many models, assessed by deviance difference test $(p>0.05)$. For those for which it was not, a non-proportional odds model was fit and results matched between the proportional odds model were found to be significant in at least one level for the non-proportional odds model. The proportional odds model was used to estimate effects for these data. As suggested by @agresti2006, if it is plausible to fit an ordinary linear regression model fitting the latent variable measured by the ordinal score response, then the assumption of proportional odds is sensible. Furthermore, the effects of interest are on the aggregate effect across all score levels of the response, which is provided by the proportional odds model. In addition, we also fit adjacent category non-proportional odds models to data that may violate the assumption. The outputs of these models are available in this supplement. While these models by design provide estimates for each transition between ordinal levels, results were highly comparable. For clarity and consistency, the results from proportional odds model fits from `VGAM::vlgm` are used in the figures of the manuscript.

```{r load-packages, warning=FALSE, message=FALSE}
# install packages if not already installed
# if(!require(tidyverse)) install.packages("tidyverse")
# if(!require(tidyr)) install.packages("tidyr")
# if(!require(VGAM)) install.packages("VGAM")
# if(!require(texreg)) install.packages("texreg")
# if(!require(readxl)) install.packages("readxl")
# if(!require(here)) install.packages("here")
# if(!require(purrr)) install.packages("purrr")
# if(!require(VGAM)) install.packages("VGAM")
# if(!require(gt)) install.packages("gt")
# if(!require(report)) install.packages("report")

# packages used in this analysis.
# library(tidyverse, quietly = TRUE) # for tidy functions
library(dplyr)
library(here)   # for file paths (optional)
library(tidyr)  # additional tidy tools
library(purrr)  # additional tidy tools
library(VGAM)   # the package that fits Proportional Odds Cumulative Logit Model
library(texreg) # optional: for plot tables
library(readxl) # for reading excel files
library(gt) # for tables
library(report) # for reporting packages and env.

```

```{r, echo=FALSE, message=FALSE}
# load tidy data & ensure column types
data_fig1 <-
  readxl::read_excel(here("data/S4_Table.xlsx"),
                     sheet = "Figure 1") %>%
  mutate(
    score = ordered(factor(score)),
    genotype = relevel(factor(genotype),"WT"),
    pretreatment = relevel(factor(pretreatment), "Mock"),
    strain = relevel(factor(strain), "S288c")
  )
  

data_fig5 <- readxl::read_excel(here("data/S4_Table.xlsx"),
                                sheet = "Figure 5") %>%
  mutate(
    score = droplevels(ordered(factor(score))),
    genotype = relevel(factor(genotype),"WT"),
    pretreatment = relevel(factor(pretreatment), "Mock"),
    strain = relevel(factor(strain), "YPS606")
  )

data_fig8 <- readxl::read_excel(here("data/S4_Table.xlsx"),
                                sheet = "Figure 8") %>%
  mutate(
    score = droplevels(ordered(factor(score))),
    genotype = relevel(factor(genotype),"ctt1∆"),
    pretreatment = relevel(factor(pretreatment), "Mock"),
    strain = relevel(factor(strain), "M22")
  )

```

```{r}
# custom functions

# pvalue formatting
my_pvalue_fun = function(x) {
    if_else(
      is.na(x), 
      NA_character_,
      if_else(x < 0.001, format(x, digits = 3, scientific = TRUE), format(round(x, 3), scientific = F))
    )
}

# Define the function to fit the model and create a summary table, accepts already filtered data
fit_and_summarize <- function(filtered_data, link_function = "cumulative") {
  model <- vglm(score ~ genotype + stress_conc,
                   family=VGAM::cumulative(parallel=TRUE),
                   data = filtered_data)
  
  table <- summary(model)@coef3
  
  list(model = model, table = table)
}

merge_and_stack_tables <- function(strain_results, strain) {
  etoh_tables <- lapply(strain_results$EtOH, `[[`, "table")
  nacl_tables <- lapply(strain_results$NaCl, `[[`, "table")

  tbl_stack(c(etoh_tables, nacl_tables),
            group_header = c(paste0(strain," EtOH: ", names(etoh_tables)),
                             paste0(strain," NaCl: ", names(nacl_tables)))
  )
}


# fit non-proportional odds models
fit_non_prop <- function(filtered_data, link_function = "logit") {
  model <- try(vglm(score ~ genotype + stress_conc,
       family=VGAM::cumulative(parallel=FALSE),
                   data = filtered_data))
  
  if(class(model) == "try-error") {
    model <- try(vglm(score ~ genotype + stress_conc,
       family=VGAM::cumulative(parallel=FALSE~stress_conc
                               ),
                   data = filtered_data))
  }
  
    table <- summary(model)@coef3
  
    list(model = model, table = table)
}


# Function to manually extract summary from vglm
extract_vglm_summary <- function(model, model_name) {
  coef_info <- summary(model)  # Get the summary of the model
  coefficients <- coef(summary(model))  # Extract coefficients and statistics
  
  # Create a data frame
  tidy_summary <- data.frame(
    term = rownames(coefficients),
    estimate = coefficients[, "Estimate"],
    std.error = coefficients[, "Std. Error"],
    statistic = coefficients[, "z value"],
    p.value = coefficients[, "Pr(>|z|)"],
    model = model_name
  )
  
  # Filter for terms containing 'genotype'
  tidy_summary <- tidy_summary[grep("genotype|pretreatment", tidy_summary$term), ]
  
  return(tidy_summary)
}

# Function to calculate Deviance Difference p-value
dev_diff_test <- function(model1, model2) {
  pchisq(
  (max(
    deviance(model1),
    deviance(model2)
  ) - min(
    deviance(model1),
    deviance(model2)
  ))
  ,
  df =
    (max(
      df.residual(model1),
      df.residual(model2)
    ) -
    min(
      df.residual(model1),
      df.residual(model2)
    )),
  lower.tail = FALSE
) 
}


```

```{r}
fit.add.op.s288c.ctt1.EtOH.prestressOnly <- data_fig1 %>%
  filter(strain == "S288c",
    pretreatment == "EtOH") %>%
  fit_and_summarize()
```

```{r}
# Only comparison where genotype is constant, formula written manually.
fit.add.op.YPS163.ctt1.NaCl <- data_fig1 %>%
  filter(strain == "YPS163",
         pretreatment != "EtOH",
         genotype == "ctt1∆") %>%
  vglm(score ~ pretreatment + stress_conc,
       family=VGAM::cumulative(parallel=TRUE),
                   data = .)
```

```{r}
# List of genotype groups
genotype_groups_fig5 <- list(
  msn24.v.WT = c("WT", "msn2∆ msn4∆"),
  skn7.v.WT = c("WT", "skn7∆"),
  ctt1yap1.v.ctt1 = c("ctt1∆", "yap1∆ ctt1∆"),
  ctt1skn7.v.ctt1 = c("ctt1∆", "skn7∆ ctt1∆"),
  ctt1msn24.v.ctt1 = c("ctt1∆", "msn2∆ msn4∆ ctt1∆")
)

# Define pretreatments
pretreatments_fig5 <- c("EtOH", "NaCl")

# Define strains
strains_fig5 <- c("YPS606")

# initialize results list
results_fig5 <- list()

# Nested looping through each strain, each pretreatment and each genotype group
for (strain_i in strains_fig5) {
  results_fig5[[strain_i]] <- list()
  for (pretreatment_i in pretreatments_fig5) {
    results_fig5[[strain_i]][[pretreatment_i]] <- list()
    for (genotype_i in names(genotype_groups_fig5)) {
      genotypes_fig5 <- genotype_groups_fig5[[genotype_i]]
      
      filtered_data <- data_fig5 %>%
        filter(strain == strain_i, pretreatment == pretreatment_i, genotype %in% genotypes_fig5)

      results_fig5[[strain_i]][[pretreatment_i]][[genotype_i]] <- fit_and_summarize(filtered_data)
    }
  }
}

```

```{r}
genotype_groups_fig8 <- list(
  ctt1gsh1.v.ctt1 = c("ctt1∆", "ctt1∆ gsh1∆")
)

strains_fig8 <- c("YPS606", "YPS163", "M22")

# Define pretreatments
pretreatments_fig8 <- c("EtOH", "NaCl")

# initialize results list
results_fig8 <- list()

# Nested looping through each strain, each pretreatment and each genotype group
for (strain_i in strains_fig8) {
  results_fig8[[strain_i]] <- list()
  for (pretreatment_i in pretreatments_fig8) {
    results_fig8[[strain_i]][[pretreatment_i]] <- list()
    for (genotype_i in names(genotype_groups_fig8)) {
      genotypes_fig8 <- genotype_groups_fig8[[genotype_i]]
      
      filtered_data <- data_fig8 %>%
        filter(strain == strain_i, pretreatment == pretreatment_i, genotype %in% genotypes_fig8)

      results_fig8[[strain_i]][[pretreatment_i]][[genotype_i]] <- fit_and_summarize(filtered_data)
    }
  }
}
```

# Estimated Coefficients of Interest

```{r, echo=FALSE}
#include=FALSE
figure_1_models = list(
  "S288c;EtOH;WTvsCtt1" = fit.add.op.s288c.ctt1.EtOH.prestressOnly$model,
  "YPS163;NaClvsMock;Ctt1" = fit.add.op.YPS163.ctt1.NaCl
)

figure_5_models = list(
  `YPS606;EtOH;WTvsMsn24`      = results_fig5$YPS606$EtOH$msn24.v.WT$model,
  `YPS606;EtOH;WTvsSkn7`       = results_fig5$YPS606$EtOH$skn7.v.WT$model,
  `YPS606;EtOH;Ctt1vsCtt1Yap1` = results_fig5$YPS606$EtOH$ctt1yap1.v.ctt1$model,
  `YPS606;EtOH;Ctt1vsCtt1Skn7` = results_fig5$YPS606$EtOH$ctt1skn7.v.ctt1$model,
  `YPS606;NaCl;Ctt1vsCtt1Yap1` = results_fig5$YPS606$NaCl$ctt1yap1.v.ctt1$model,
  `YPS606;NaCl;Ctt1vsCtt1Skn7` = results_fig5$YPS606$NaCl$ctt1skn7.v.ctt1$model
)

figure_8_models = list(
  "YPS606;EtOH;Ctt1vsCtt1Gsh1" = results_fig8$YPS606$EtOH$ctt1gsh1.v.ctt1$model,
  "YPS163;EtOH;Ctt1vsCtt1Gsh1" = results_fig8$YPS163$EtOH$ctt1gsh1.v.ctt1$model,
  "M22;EtOH;Ctt1vsCtt1Gsh1" = results_fig8$M22$EtOH$ctt1gsh1.v.ctt1$model,
  "YPS606;NaCl;Ctt1vsCtt1Gsh1" = results_fig8$YPS606$NaCl$ctt1gsh1.v.ctt1$model,
  "YPS163;NaCl;Ctt1vsCtt1Gsh1" = results_fig8$YPS163$NaCl$ctt1gsh1.v.ctt1$model,
  "M22;NaCl;Ctt1vsCtt1Gsh1" = results_fig8$M22$NaCl$ctt1gsh1.v.ctt1$model
)
```

```{r, echo=FALSE}
# include=FALSE,
# summarise model's estimates of interest
model_summaries_fig1 <- map2_df(figure_1_models, names(figure_1_models), extract_vglm_summary) %>%
  mutate(adj.p.value = p.adjust(p = p.value, n=14, method="fdr") %>% my_pvalue_fun(.),
         p.value = my_pvalue_fun(as.numeric(p.value))) %>%
  tidyr::separate(model, into = c("Strain", "Treatment", "Genotype"), sep = ";") %>%
  select(Strain, Treatment, Genotype, estimate,std.error, statistic, p.value, adj.p.value)

model_summaries_fig5 <- map2_df(figure_5_models, names(figure_5_models), extract_vglm_summary) %>%
  mutate(adj.p.value = p.adjust(p = p.value, n=14, method="fdr") %>% my_pvalue_fun(.),
         p.value = my_pvalue_fun(as.numeric(p.value))) %>%
  tidyr::separate(model, into = c("Strain", "Treatment", "Genotype"), sep = ";") %>%
  select(Strain, Treatment, Genotype, estimate,std.error, statistic, p.value, adj.p.value)

model_summaries_fig8 <- map2_df(figure_8_models, names(figure_8_models), extract_vglm_summary) %>%
  mutate(adj.p.value = p.adjust(p = p.value, n=14, method="fdr") %>% my_pvalue_fun(.),
         p.value = my_pvalue_fun(as.numeric(p.value))) %>%
  tidyr::separate(model, into = c("Strain", "Treatment", "Genotype"), sep = ";") %>%
  select(Strain, Treatment, Genotype,  estimate,std.error, statistic, p.value, adj.p.value) 
```

```{r}
library(gt)

# Create gt tables
gt_table_fig1 <- gt(model_summaries_fig1) %>% 
  tab_header(title = "Fig 1:  Noted Comparisons") %>%
  cols_label(
    estimate = "Estimate",
    p.value = "P-Value",
    adj.p.value = "FDR",
    Genotype = "Comparison"
  ) %>%
  fmt_number(
    columns = c(estimate, std.error, statistic, p.value), n_sigfig = 3
    #decimals = 4
  ) %>%
  fmt_scientific(
    columns = p.value,
    n_sigfig =3
  ) %>%
  cols_align(
    align = "center",
    columns = everything()
  ) %>%
  tab_footnote(
    footnote = "FDR corresponds to Benjamini & Hochberg correction method",
    locations = cells_column_labels(adj.p.value)
  )

gt_table_fig5 <- gt(model_summaries_fig5) %>%
  tab_header(title = "Fig 5: Noted Comparisons") %>%
  cols_label(
    estimate = "Estimate",
    p.value = "P-Value",
    adj.p.value = "FDR",
    Genotype = "Comparison"
  ) %>%
  fmt_number(
    columns = c(estimate, std.error, statistic, p.value),n_sigfig = 3
    #decimals = 4
  ) %>%
  fmt_scientific(
    columns = p.value,
    n_sigfig =3
  ) %>%
  cols_align(
    align = "center",
    columns = everything()
  ) #%>%
  #tab_footnote(
  #  footnote = "FDR corresponds to Benjamini & Hochberg correction method",
  #  locations = cells_column_labels(adj.p.value)
  #)

gt_table_fig8 <- gt(model_summaries_fig8) %>%
  tab_header(title = "Fig 8: Noted Comparisons") %>%
  cols_label(
    estimate = "Estimate",
    p.value = "P-Value",
    adj.p.value = "FDR",
    Genotype = "Comparison"
  ) %>%
  fmt_scientific(
    columns = p.value,
    n_sigfig =3
  ) %>%
  fmt_number(
    columns = c(estimate, std.error, statistic, p.value),n_sigfig = 3
  ) %>%
  cols_align(
    align = "center",
    columns = everything()
  ) #%>%
  #tab_footnote(
  #  footnote = "FDR corresponds to Benjamini & Hochberg correction method",
  #  locations = cells_column_labels(adj.p.value)
  #)

```

```{r, results='asis', echo=FALSE}
# Print the tables
gt_table_fig1
gt_table_fig5
# gt_table_fig8 
```

\newpage

```{r, results='asis', echo=FALSE}
# Print the tables
gt_table_fig8 
```

# Testing Proportional Odds Assumption

```{r, warning=FALSE, message=FALSE, error=FALSE}
# Check for proportional odds assumption Fig 1
fit.add.o.s288c.ctt1.EtOH.prestressOnly <- data_fig1 %>%
  filter(strain == "S288c",
    pretreatment == "EtOH") %>%
  vglm(score ~ genotype + stress_conc,
       family=cumulative(parallel=FALSE),
                   data = .)

dev_diff_pval_fig1_s288c <-
anova(fit.add.o.s288c.ctt1.EtOH.prestressOnly, fit.add.op.s288c.ctt1.EtOH.prestressOnly$model, type = "I")$`Pr(>Chi)`[[2]] %>% round(3)

fit.add.o.YPS163.ctt1.NaCl <- data_fig1 %>%
  filter(strain == "YPS163",
         pretreatment != "EtOH",
         genotype == "ctt1∆") %>%
  vglm(score ~ pretreatment + stress_conc,
       family=cumulative(parallel=FALSE),
                   data = .)

## see these are comparable
dev_diff_pval_fig1_YPS163_alt <- pchisq(
  deviance(fit.add.op.YPS163.ctt1.NaCl) -
    deviance(fit.add.o.YPS163.ctt1.NaCl),
       df = df.residual(fit.add.op.YPS163.ctt1.NaCl) -
    df.residual(fit.add.o.YPS163.ctt1.NaCl),
  lower.tail = FALSE)

dev_diff_pval_fig1_YPS163 <- anova(fit.add.o.YPS163.ctt1.NaCl, fit.add.op.YPS163.ctt1.NaCl, type = "I")$`Pr(>Chi)`[[2]] %>% round(3)
```

```{r, warning=FALSE, message=FALSE,error=FALSE,include=FALSE}
# , include=FALSE
results_fig5_non_prop <- list()

# Nested looping through each strain, each pretreatment and each genotype group
for (strain_i in strains_fig5) {
  results_fig5_non_prop[[strain_i]] <- list()
  for (pretreatment_i in pretreatments_fig5) {
    results_fig5_non_prop[[strain_i]][[pretreatment_i]] <- list()
    for (genotype_i in names(genotype_groups_fig5)) {
      genotypes_fig5 <- genotype_groups_fig5[[genotype_i]]
      
      filtered_data <- data_fig5 %>%
        filter(strain == strain_i, pretreatment == pretreatment_i, genotype %in% genotypes_fig5)

      results_fig5_non_prop[[strain_i]][[pretreatment_i]][[genotype_i]] <- try(fit_non_prop(filtered_data))
    }
  }
}
```

The null hypothesis for this deviance difference test suggest that the proportional odds assumption is consistent with the data. The assumption is supported in some comparisons and inconsistent with the data in others. However, the underlying logic of a continuous latent variable being approximated by the semi-quantitative scores is reasonable and more easily interpreted. As such, values reported here are for models with proportional odds.

```{r, warning=FALSE}
dev_diff_pval_fig5_YPS606_msn24vWT_et <- 
  anova(results_fig5_non_prop$YPS606$EtOH$msn24.v.WT$model, results_fig5$YPS606$EtOH$msn24.v.WT$model, type = "I")$`Pr(>Chi)`[[2]] %>% round(3)

dev_diff_pval_fig5_YPS606_skn7vWT_et <-
anova(results_fig5_non_prop$YPS606$EtOH$skn7.v.WT$model, results_fig5$YPS606$EtOH$skn7.v.WT$model, type = "I")$`Pr(>Chi)`[[2]] %>% round(3)

dev_diff_pval_fig5_YPS606_ctt1skn7vctt1_et <-
anova(results_fig5_non_prop$YPS606$EtOH$ctt1skn7.v.ctt1$model, results_fig5$YPS606$EtOH$ctt1skn7.v.ctt1$model, type = "I")$`Pr(>Chi)`[[2]] %>% round(3)

dev_diff_pval_fig5_YPS606_ctt1yap1vctt1_et <-
anova(results_fig5_non_prop$YPS606$EtOH$ctt1yap1.v.ctt1$model, results_fig5$YPS606$EtOH$ctt1yap1.v.ctt1$model, type = "I")$`Pr(>Chi)`[[2]] %>% round(3)

dev_diff_pval_fig5_YPS606_ctt1skn7vctt1_nacl <-
anova(results_fig5_non_prop$YPS606$NaCl$ctt1skn7.v.ctt1$model, results_fig5$YPS606$NaCl$ctt1skn7.v.ctt1$model, type = "I")$`Pr(>Chi)`[[2]] %>% round(3)

dev_diff_pval_fig5_YPS606_ctt1yap1vctt1_nacl <-
anova(results_fig5_non_prop$YPS606$NaCl$ctt1yap1.v.ctt1$model, results_fig5$YPS606$NaCl$ctt1yap1.v.ctt1$model, type = "I")$`Pr(>Chi)`[[2]] %>% round(3)
```

```{r, warning=FALSE, message=FALSE}
results_fig8_non_prop <- list()

for (strain_i in strains_fig8) {
  results_fig8_non_prop[[strain_i]] <- list()
  for (pretreatment_i in pretreatments_fig8) {
    results_fig8_non_prop[[strain_i]][[pretreatment_i]] <- list()
    for (genotype_i in names(genotype_groups_fig8)) {
      genotypes_fig8 <- genotype_groups_fig8[[genotype_i]]
      
      filtered_data <- data_fig8 %>%
        filter(strain == strain_i, pretreatment == pretreatment_i, genotype %in% genotypes_fig8)

      results_fig8_non_prop[[strain_i]][[pretreatment_i]][[genotype_i]] <- try(fit_non_prop(filtered_data))
    }
  }
}
```

```{r, warning=FALSE}
dev_diff_pval_fig8_YPS606_ctt1gsh1vctt1_nacl <- dev_diff_test(
  results_fig8_non_prop$YPS606$NaCl$ctt1gsh1.v.ctt1$model,
  results_fig8$YPS606$NaCl$ctt1gsh1.v.ctt1$model) %>%
  signif(2)
# anova(results_fig8_non_prop$YPS606$NaCl$ctt1gsh1.v.ctt1$model, results_fig8$YPS606$EtOH$ctt1gsh1.v.ctt1$model, type = "I")$`Pr(>Chi)`[[2]] 


dev_diff_pval_fig8_YPS163_ctt1gsh1vctt1_nacl <- dev_diff_test(
  results_fig8_non_prop$YPS163$NaCl$ctt1gsh1.v.ctt1$model,
  results_fig8$YPS163$NaCl$ctt1gsh1.v.ctt1$model) %>%
  signif(2)
# anova(results_fig8_non_prop$YPS163$NaCl$ctt1gsh1.v.ctt1$model, results_fig8$YPS163$NaCl$ctt1gsh1.v.ctt1$model, type = "I")$`Pr(>Chi)`[[2]] %>% round(3)

dev_diff_pval_fig8_M22_ctt1gsh1vctt1_nacl <- dev_diff_test(
  results_fig8_non_prop$M22$NaCl$ctt1gsh1.v.ctt1$model,
  results_fig8$M22$NaCl$ctt1gsh1.v.ctt1$model) %>% 
  signif(2)
# anova(results_fig8_non_prop$M22$NaCl$ctt1gsh1.v.ctt1$model, results_fig8$M22$NaCl$ctt1gsh1.v.ctt1$model, type = "I")$`Pr(>Chi)`[[2]] %>% round(3)

dev_diff_pval_fig8_YPS606_ctt1gsh1vctt1_et <- dev_diff_test(
  results_fig8_non_prop$YPS606$EtOH$ctt1gsh1.v.ctt1$model,
  results_fig8$YPS606$EtOH$ctt1gsh1.v.ctt1$model) %>%
  signif(2)
# anova(results_fig8_non_prop$YPS606$EtOH$ctt1gsh1.v.ctt1$model, results_fig8$YPS606$EtOH$ctt1gsh1.v.ctt1$model, type = "I")$`Pr(>Chi)`[[2]] %>% round(3)

dev_diff_pval_fig8_YPS163_ctt1gsh1vctt1_et <- dev_diff_test(
  results_fig8_non_prop$YPS163$EtOH$ctt1gsh1.v.ctt1$model,
  results_fig8$YPS163$EtOH$ctt1gsh1.v.ctt1$model) %>%
  signif(2)
# anova(results_fig8_non_prop$YPS163$EtOH$ctt1gsh1.v.ctt1$model, results_fig8$YPS163$EtOH$ctt1gsh1.v.ctt1$model, type = "I")$`Pr(>Chi)`[[2]] %>% round(3)

dev_diff_pval_fig8_M22_ctt1gsh1vctt1_et <- dev_diff_test(
  results_fig8_non_prop$M22$EtOH$ctt1gsh1.v.ctt1$model,
  results_fig8$M22$EtOH$ctt1gsh1.v.ctt1$model) %>%
  signif(2)
# anova(results_fig8_non_prop$M22$EtOH$ctt1gsh1.v.ctt1$model, results_fig8$M22$EtOH$ctt1gsh1.v.ctt1$model, type = "I")$`Pr(>Chi)`[[2]] %>% round(3)
```

```{r}
prop.testing.results <-
  data.frame(
    Figure = c(1,1,rep(5,6), rep(8,6)),
    Strain = c(
      "S288c",
      "YPS163",
      rep("YPS606",6),
      rep(c("YPS163", "YPS606", "M22"),2)
    ),
    Pretreatment = c(
      "EtOH",
      "NaCl vs Mock",
      rep("EtOH", 4),
      rep("NaCl",5),
      rep("EtOH",3)
    ),
    Genotype =
      c(
        "WT vs ctt1∆",
        "ctt1∆",
        "msn2msn4∆∆ vs WT",
        "skn7∆ vs WT",
        "ctt1∆skn7∆ vs ctt1∆",
        "ctt1∆yap1∆ vs ctt1∆",
        "ctt1∆skn7∆ vs ctt1∆",
        "ctt1∆yap1∆ vs ctt1∆",
        rep("ctt1∆gsh1∆ vs ctt1∆",6)
      ),
    P.Value = 
      c(
        dev_diff_pval_fig1_s288c,
        dev_diff_pval_fig1_YPS163,
        dev_diff_pval_fig5_YPS606_msn24vWT_et,
        dev_diff_pval_fig5_YPS606_skn7vWT_et,
        dev_diff_pval_fig5_YPS606_ctt1skn7vctt1_et,
        dev_diff_pval_fig5_YPS606_ctt1yap1vctt1_et,
        dev_diff_pval_fig5_YPS606_ctt1skn7vctt1_nacl,
        dev_diff_pval_fig5_YPS606_ctt1yap1vctt1_nacl,
        dev_diff_pval_fig8_YPS163_ctt1gsh1vctt1_et,
        dev_diff_pval_fig8_YPS606_ctt1gsh1vctt1_et,
        dev_diff_pval_fig8_M22_ctt1gsh1vctt1_et,
        dev_diff_pval_fig8_YPS163_ctt1gsh1vctt1_nacl,
        dev_diff_pval_fig8_YPS606_ctt1gsh1vctt1_nacl,
        dev_diff_pval_fig8_M22_ctt1gsh1vctt1_nacl
      )
  )

prop.testing.results %>% gt() %>%
  tab_header(title = "Tests of Proportional Odds Assumption") %>%
  cols_label(
    # term = "Estimator",
    # estimate = "Estimate",
    P.Value = "P-Value"#,
    # Genotype = "Comparison"
    # model = "Model"
  ) %>%
  fmt_scientific(
    columns = P.Value,
    rows = P.Value < 0.001,
    drop_trailing_zeros = TRUE,
    n_sigfig = 2
  ) %>%
  # fmt_number(
  #   columns = c(estimate, std.error, statistic, p.value),n_sigfig = 3
  #   #decimals = 6
  # ) %>%
  cols_align(
    align = "center",
    columns = everything()
  )
```

## Ensuring significance without proportional odds assumption

For models for which there was evidence of non-proportional odds, a cumulative logit model without a proportional odds assumption was fit and estimates for the contrast of interest were extracted. In general, the system of equations for the non-proportional odds is:

$$
\begin{split}
log{\frac{P(Y\le j)}{P(Y>j)}} &= \alpha_j + \beta_{j, gene\Delta}X_{gene\Delta} + \beta_{j,[H_2O_2]}X_{[H_2O_2]} \\
log{\frac{P(Y\le j)}{P(Y>j)}} &= \alpha_j + \beta_{j, prestress}X_{prestress} + \beta_{j, [H_2O_2]}X_{[H_2O_2]}
\end{split}
$$

Note that the effects of $X$ values are allowed to differ for different logits in this model. To illustrate, the system of equations for the cumulative logit model comparing Figure 1's S288c during EtOH pretreatment comparing $WT$ vs $ctt1\Delta$, with the proportional odds model:

$$
\begin{split}
log{\frac{P(Y\le 0)}{P(Y>0)}} &= \alpha_0 + \beta_{ gene\Delta}X_{gene\Delta} + \beta_{[H_2O_2]}X_{[H_2O_2]} \\
&= -7.49 -6.68 X_{ctt1\Delta} + 8.36X_{[H_2O_2]}\\
log{\frac{P(Y\le 1)}{P(Y=2)}} &= \alpha_1 + \beta_{ gene\Delta}X_{gene\Delta} + \beta_{[H_2O_2]}X_{[H_2O_2]} \\
&= -5.04 -6.68 X_{ctt1\Delta} + 8.36X_{[H_2O_2]}\\
\end{split}
$$

and the non-proportional odds model:
$$
\begin{split}
    log{\frac{P(Y\le 0)}{P(Y>0)}} &= \alpha_0 + \beta_{\mathbf{0}, gene\Delta}X_{gene\Delta} + \beta_{\mathbf{0},[H_2O_2]}X_{[H_2O_2]} \\ 
    &= -6.70 -6.45 X_{ctt1\Delta} + 7.67X_{[H_2O_2]}\\
    log{\frac{P(Y\le 1)}{P(Y=2)}} &= \alpha_1 + \beta_{\mathbf{1}, gene\Delta}X_{gene\Delta} + \beta_{\mathbf{1},[H_2O_2]}X_{[H_2O_2]}  \\
    &= -5.61 -6.10 X_{ctt1\Delta} + 8.47X_{[H_2O_2]}
\end{split}
$$
Note that the estimated effects for genotype and hydrogen peroxide can vary between each logit in the system of equations, with positive coefficients, as seen here for hydrogen peroxide, corresponding to an increased probability of the sample belonging to a lower score as that covariate increases. 
The estimates for $\beta_j$ for all comparisons for which the proportional odds assumption is statistically violated are plotted below. Red corresponds to a 95% confidence interval (CI) that does not contain 0 and blue to non-significance. Note that the only comparison for which no estimate is nonzero is the YPS606 comparison from Figure 8A, which was also non-significant in the proportional-odds model estimate shown above. This line of evidence supports the robustness of our statistical interpretations.

```{r, fig.width=6, fig.height=9,dpi=600, message=FALSE}
plotreg(list(
             fit.add.o.YPS163.ctt1.NaCl,
             results_fig5_non_prop$YPS606$NaCl$ctt1skn7.v.ctt1$model,
             results_fig5_non_prop$YPS606$NaCl$ctt1yap1.v.ctt1$model,
             # results_fig8_non_prop$YPS606$EtOH$ctt1gsh1.v.ctt1$model,
             # results_fig8_non_prop$YPS606$NaCl$ctt1gsh1.v.ctt1$model,
             results_fig8_non_prop$YPS163$EtOH$ctt1gsh1.v.ctt1$model,
             results_fig8_non_prop$YPS163$NaCl$ctt1gsh1.v.ctt1$model,
             results_fig8_non_prop$M22$EtOH$ctt1gsh1.v.ctt1$model,
             results_fig8_non_prop$M22$NaCl$ctt1gsh1.v.ctt1$model
             ), 
        omit.coef = "(Intercept)|(stress_conc)",
          custom.model.names = c("Fig1:YPS163", 
                                 "Fig5:NaCl;skn7ctt1",
                                 "Fig5:NaCl;yap1ctt1",
                                 # "Fig8:YPS606;EtOH",
                                 # "Fig8:YPS606;NaCl",
                                 "Fig8:YPS163;EtOH",
                                 "Fig8:YPS163;NaCl",
                                 "Fig8:M22;EtOH",
                                 "Fig8:M22;NaCl"),
         lwd.vbars = 0, ci.force=TRUE,
        custom.note = "Bars denote CIs (95%) computed from SEs. \nCircle points denote significance.",
        custom.title = "Non-Proportional Odds Effect Estimates"
        )

```


```{r, results='asis'}
report(sessionInfo())
```

\newpage
